# High light can alleviate chilling stress in maize

**DOI:** 10.1101/2023.03.07.531599

**Authors:** Lee Cackett, Angela C. Burnett, Jessica Royles, Julian M. Hibberd, Johannes Kromdijk

## Abstract

Chilling stress has the potential to significantly decrease growth and yield of sensitive crop plants such as maize. Based on previous work, high light during chilling may exacerbate stress via enhanced photoinhibition but may also aid acclimation responses to chilling. To further understand molecular processes behind responses to chilling with and without high light, two maize accessions with contrasting tolerance (B73 and F7) were exposed to three treatments: chilling, chilling combined with high light and high light alone. Transcriptome data indicated that the chilling treatment resulted in the largest stress response. Addition of high light to chilling stress had a mitigating, rather than additive effect on stress, as evident from alleviated repression of photosynthesis-related genes and less induction of stress-related pathways such as abscisic acid signalling and senescence compared with the response to chilling alone. Five transcription factors belonging to well-known stress-related transcription factor families were identified as candidates for driving the transcriptional changes behind the high-light induced mitigation of chilling stress. Physiological measurements of non-photochemical quenching and the maximum quantum efficiency of photosystem II corroborated the transcriptome results, showing that the addition of high light alleviated photoinhibition and membrane damage caused by chilling. High light alone had little effect on the plant transcriptome or physiological response. Overall, this study overturns previous reports, offers a new outlook on the impact of high light during chilling stress and has the potential to provide clearer targets for crop engineering.

## Introduction

The world’s population is rising dramatically and with it the need for increased global food production (FAO, 2018). Adding to this challenge is the increasing frequency and severity of extreme weather events (IPCC, 2018) meaning that crop production must also become more climate-resilient if we are to meet the food security challenges of the future. Specifically, temperature is an important determinant of photosynthesis and crop yield (Yoshida et al., 1981; Geange et al., 2021). Therefore, in order to maintain crop yield stability in future environments it is imperative that we continue to develop crop plants which are increasingly resilient to temperature extremes and are adapted to growing and thriving in different climatic conditions. To meet this challenge, a more extensive understanding of the physiological and molecular foundations of temperature stress responses is required.

Maize is the most-produced cereal crop in the world, with a production mass of 1.1 billion tonnes in 2019 (FAO, 2021). It has diverse uses including food, feed, energy, and raw materials for industry (Ma et al., 2022) and makes a major contribution to the caloric intake of the global population (Erenstein et al., 2022). Maize is highly diversified at a genomic level and has dispersed into a wide range of environments distinct from its environment of domestication (Tenaillon and Charcosset, 2011; Sowiński et al., 2020). Despite this diversification, the tropical origins of maize mean that it is highly susceptible to chilling stress when grown in temperate locations. Here, chilling is defined as suboptimal cool temperatures below 15°C and above 0°C (i.e. not freezing stress) with severe chilling stress occurring below 8°C (Frascaroli and Revilla, 2019; Burnett and Kromdijk, 2022). Chilling stress significantly limits seed germination, seedling establishment, and subsequent plant growth and development including yield (Jompuk et al., 2005; Revilla et al., 2005; Sanghera et al., 2011). For example, cold snaps – short periods of chilling temperatures – result in significant yield losses (Sanghera et al., 2011). In addition, poor early-season establishment can elevate farm costs associated with suppression of weeds via herbicide applications or mechanical measures.

The morphological and physiological impacts of chilling stress on maize have been extensively studied and recently reviewed (Burnett and Kromdijk 2022; Ma et al., 2022). Chilling stress in maize results in excessive wilting, chlorosis and necrosis and decreased germination rate, root growth, leaf area and leaf appearance rate (Revilla et al., 2005; Frascaroli and Revilla 2019). Physiologically, chilling stress in maize results in increased cell membrane disruption (Miedema, 1982), decreased chlorophyll content and decreased operating efficiency of photosystem II (Φ_PSII_) (reviewed by Burnett and Kromdijk, 2022). Further, chilling stress in maize decreases the maximum quantum efficiency of photosystem II (F_v_/F_m_) (Dolstra et al., 1994, Fracheboud et al., 1999, Ying et al., 2002) and increases non-photochemical quenching (NPQ) (Fracheboud et al., 1999, Rodríguez et al., 2013, Riva-Roveda et al., 2016). NPQ comprises rapidly-reversible photoprotective responses as well as damage-related photoinhibitive processes from which the plant requires more time to recover (Baker, 2008, Malnoë, 2018).

These physiological changes resulting from chilling stress contribute to lowered rates of photosynthesis, ultimately impacting growth, morphology, and productivity, making chilling tolerance of maize an important target for breeding (Burnett and Kromdijk, 2022). Many transcription factors (TFs), alleles and quantitative trait loci (QTL) related to chilling stress in maize have been identified (Fracheboud et al., 2004; Waters et al., 2017; Zeng et al., 2021; Ma et al., 2022). For example, genes such as *CYTOKININ RESPONSE REGULATOR 1 (CRR1), MAP KINASE 8* (*MPK8)* and the *DEHYDRATION-RESPONSIVE ELEMENT-BINDING proteins (DREBs)* have been shown to influence chilling tolerance of maize varieties (Qin et al., 2004, Wang and Dong, 2009, Wang, Yang and Yang, 2011, Zeng et al., 2021). Despite this progress, the factors underpinning chilling tolerance in maize, and determining whether a genotype will display chilling tolerance or chilling sensitivity, are not completely understood (Frascaroli and Revilla, 2019). Notably, significant gaps remain in knowledge of the molecular foundations which govern the well-known physiological responses of maize to low temperatures, specifically key genes controlling signalling pathways and responses (reviewed in Sowiński et al., 2020).

Photosynthesis is particularly light- and temperature-sensitive due to the tight coupling between the thylakoid reactions and downstream C_4_ acid shuttle and Calvin-Benson-Bassham cycle (Hüner et al., 1998). Chilling stress and high light often coincide during clear-sky cold snaps in temperate regions. High light levels can aid maize growth but can also cause photoinhibition. In the context of maize chilling responses, the role of high light is currently unclear, with two competing mechanisms reported across literature. On the one hand, combined chilling and high light stress has long been considered as additive, with high light exacerbating chilling stress and vice versa (Ortiz-Lopez, 1990). Indeed, it has been demonstrated that high light combined with chilling stress increases photoinhibition in field-grown maize (Farage and Long, 1987). Conversely, other studies have reported that high light can aid acclimation to chilling stress and thereby mitigate the negative effects of chilling. For example, increasing light intensity during a mild chilling treatment enhanced tolerance to a subsequent severe chilling event (Szalai et al., 2018), a phenomenon known as stress priming (Liu et al., 2022). In addition, several cold-tolerant maize lines evaluated by Grzybowski et al. (2019) displayed more photo-sensitivity than cold-sensitive lines in the same study, suggesting that exposure to high light does not have to be additive with chilling stress.

In this work we explored the relationship between chilling and high light in greater depth, in order to elucidate the possible mitigation of chilling stress by high light. In particular, we used a combined genetic and physiological approach to determine whether high light plays a role in exacerbation or mitigation of stress induced by a short cold spell (a cold night and subsequent cool morning at two contrasting light levels, 400 and 1000 μmol photons m^-2^ s^-1^). We selected two contrasting maize inbred lines: a chilling tolerant accession, F7 (a European Flint line) and a mildly chilling-sensitive accession, B73 (an American Dent line); these accessions have previously been observed to have different chilling tolerances in a two to four day chilling stress period (Enders et al., 2018). Specifically, we set out to test the following hypotheses: 1. High light will exacerbate chilling stress in maize leading to greater physiological damage and reduced photosynthesis. 2. The chilling sensitive accession (B73) will be more susceptible to chilling stress than the chilling tolerant accession (F7). 3. Physiological responses to high light and chilling stress are underpinned by transcriptomic responses, and differences in physiological and photosynthetic changes will reflect differences in gene expression.

The transcriptomic responses confirmed the expected difference in chilling tolerance between F7 and B73, based on both the overall magnitude of the transcriptomic response, as well as changes in transcript abundance of previously characterised cold-responsive genes. However, surprisingly transcriptomic and physiological data in both accessions indicated a mitigating effect of high light on the chilling stress. Five candidate transcription factors were identified as control factors for the mitigating effect of high light on chilling, all of which belong to transcription factor families which have been previously implicated in abiotic stress responses.

Responses to chilling in combination with 1000 µmol m^-2^ s^-1^ consistently displayed lower values for physiological stress proxies as well as for transcriptomic responses, compared to chilling in combination with 400 µmol m^-2^ s^-1^. Overall these results clearly demonstrate that high light levels can alleviate chilling stress in maize.

## Results

### High light impacted the transcriptome signature of chilling stress

Plants of two contrasting maize accessions, B73 and F7, were exposed to one of three treatments. Each treatment was composed of twelve hours night and three hours photoperiod, following which the plants were sampled. The first treatment (referred to as chilling) exposed plants to chilling temperature of 5 °C during the night and 8 °C during the photoperiod, while keeping light intensity during the three hour photoperiod the same as during growth prior to the experiment (400 µmol m^-2^ s^-1^). The second treatment (referred to as chilling plus high light), exposed plants to the same chilling temperatures of 5/8 °C N/D, but also raised the light intensity during the three morning hours to 2.5 times the growth intensity (1000 µmol m^-2^ s^-1^). Finally, the third treatment (referred to as high light) kept temperature at 20/28 °C N/D, while raising the light intensity during the three morning hours to 1000 µmol m^-2^ s^-1^. Plants from each treatment were compared to matching control plants (0/400 µmol m^-2^ s^-1^, 20/28 °C N/D) sampled at the same time of day, one day prior to the start of each treatment. To explore the molecular basis of the responses to chilling and high light, leaf tissue from all untreated and treated plants was sampled for 3’UTR RNA sequencing (RNA-seq). Principal component analysis (PCA) on the transcripts per million (TPM) data (Fig. 1A) assessed the quality and structure of the data and demonstrated that for both B73 and F7 accessions, the chilling treated samples (blue) differed most from the control samples (grey) across the first principal component (explaining 37.3% of transcriptomic variation) whilst high light treated samples (red) appeared similar to the control samples. The chilling plus high light treated samples (yellow) were positioned in between the chilling only and high light only samples, suggesting that the addition of high light to chilled plants had a mitigating effect. The second principal component, explaining 22.8% of the transcriptomic variation, mostly depicted the difference between B73 and F7 accessions (circles and triangles respectively). When TPM data from the two accessions was separated and independent PCAs performed, the differences in treatments described in Figure 1A were conserved (Fig. 1B and C), with the chilling treated samples (blue) differing the most from the control samples, followed by the chilling and high light treated samples (yellow) and the high light treated samples (red) differing very little from the control samples. This explained 43.7% and 61.7% of the transcriptomic variation for B73 and F7 accessions respectively.

**Figure 1:**
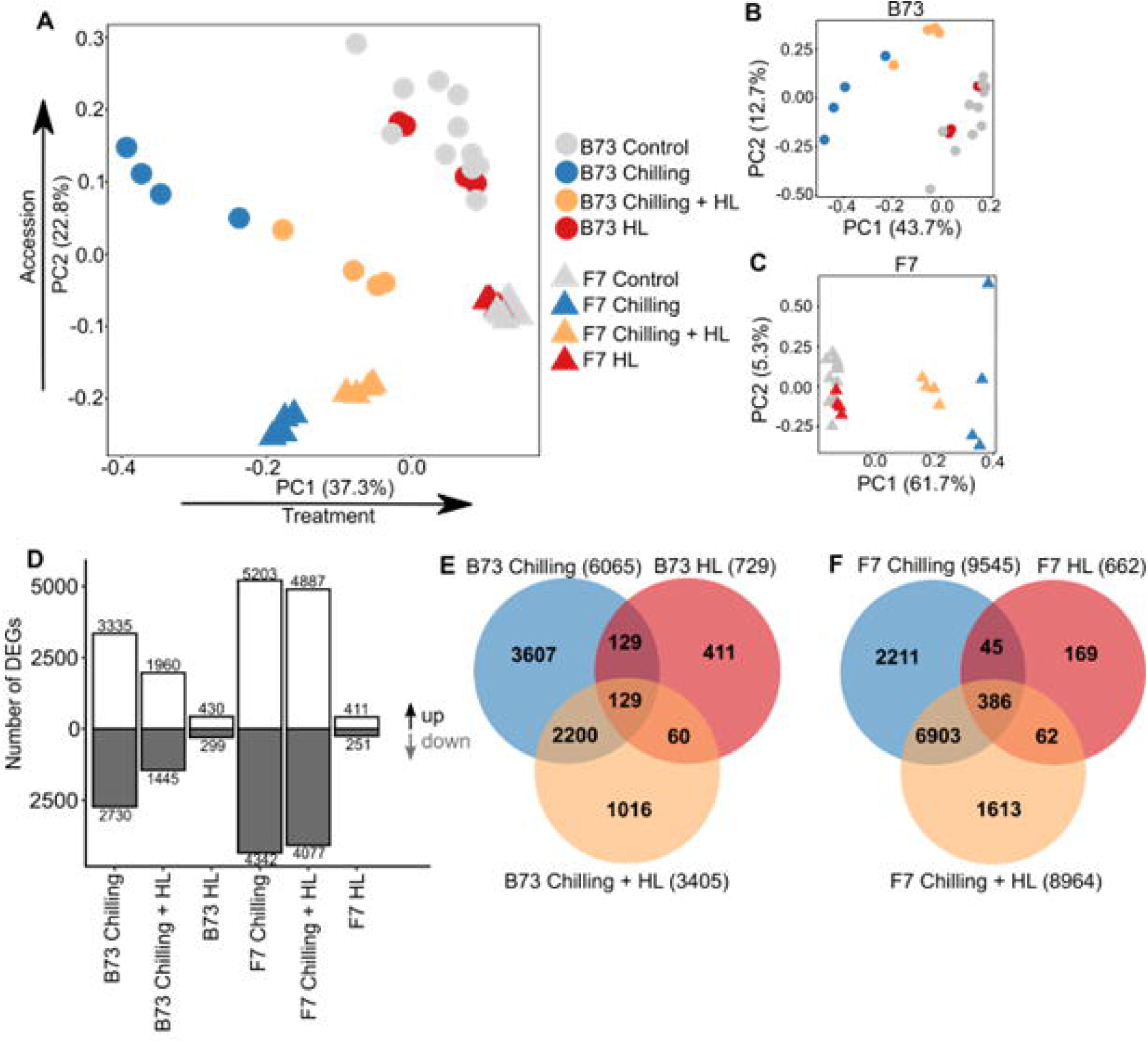
Global transcriptome responses to the chilling and high light treatments. **A**, principal component analysis (PCA) plot of all 48 samples (24 control, 8 chilling, 8 chilling plus high light (Chilling + HL), 8 high light (HL)) showing how the transcriptome data separate based on treatment and accession. **B**, PCA plot of only B73 samples (24 samples). **C**, PCA plot of only F7 samples (24 samples). For all PCA plots, transcripts per million (TPM) counts from Salmon alignments were filtered for genes with at least 36 counts across the 48 samples for **A**, and genes with at least 18 counts across the 24 samples for **B** and **C**. The filtered TPM data was then normalised using variance stabilising transformation (vst) to estimate the dispersion trend across samples. The B73 and F7 accessions are represented as a circle and triangle respectively. Chilling, chilling plus high light and high light treated samples are represented in blue, yellow, and red respectively and untreated samples in grey. **D**, number of significantly (*p*-adj. < 0.05) differently expressed genes (DEGs) in each treatment for each accession determined by a DESeq2 analysis. Up-regulated genes are represented by white bars, and down-regulated genes by grey bars. Numbers above each bar indicate the count of DEGs represented by the bar. **E**, Venn diagram showing the extent of overlap between DEGs in each treatment in B73. **F**, Venn diagram showing the extent of overlap between DEGs in each treatment in F7.

Differentially expressed genes (DEGs) for each treatment were identified by comparing the four treated samples with their four matching control samples. Only genes with statistically significant changes in expression (i.e. adjusted *p*-value < 0.05) in each treatment were retained for subsequent analyses. The chilling treatment led to differential expression of 6065 and 9545 genes for B73 and F7 respectively (Fig. 1D, E and F). Addition of high light to the chilling treatment (chilling + HL) decreased the number of DEGs in both maize lines substantially, resulting in 3405 and 8964 genes for B73 and F7 respectively. Of these genes, the majority were also differentially expressed in the chilling treatment (68% for B73 and 81% for F7, Fig. 1E and F). The transcriptome profile of the high light treatment without chilling was considerably less responsive compared with the chilling treatments, with only 729 and 662 DEGs for B73 and F7 respectively (Fig. 1D, E and F). Results of a Gene Ontology (GO) enrichment analysis on DEGs after high light treatment included the “response to high light intensity” GO term, indicating that the plants did still respond to the high light treatment, but only to a limited degree. When comparing between accessions, 73% of the chilling responsive DEGs in B73 were also differentially expressed in F7 and 70% of the chilling plus high light responsive genes in B73 were also present in the corresponding F7 list (Supp. figure 1). Only 20% of the high light responsive genes in B73 were also present in the corresponding F7. Overall, the RNA-seq data indicated that the chilling treatment alone had the greatest impact on transcription, whilst the addition of high light to chilling treatment decreased the response and the plants treated with high light showed much fewer transcriptional change.

### F7 showed transcriptional indicators of improved chilling tolerance

When comparing the transcriptome data from the two accessions, evidence for enhanced tolerance of F7 to chilling stress emerged and provided indications of the mechanisms driving such tolerance. Notably, the transcriptional response of F7 to chilling and chilling plus high light was much broader (i.e. more DEGs) compared to B73 despite using the B73 genome for alignment (Fig. 1D). This is consistent with previously observed differences in the number of cold responsive genes between tolerant and sensitive genotypes (Zhou et al., 2022a). Additionally, several individual genes which have previously been linked with chilling tolerance showed opposite patterns of expression in F7 and B73 (Fig. 2). Zeaxanthin accumulation is negatively associated with chilling tolerance in maize (Fracheboud et al., 2002). *ZEAXANTHIN EPOXIDASE 2* (*ZEP2*), which catalyses the conversion of zeaxanthin to violaxanthin, showed down-regulation in B73 in response to chilling and chilling plus high light, whereas expression was up-regulated in F7 in the treatments (Fig. 2A). Conversely, *VIOLAXANTHIN DE-EPOXIDASE 3b* (*VDE3b*) which catalyses the conversion of violaxanthin to zeaxanthin was up-regulated in B73 but down-regulated in F7 in response to chilling and chilling plus high light (Fig. 2B). Both DEGs point towards a greater accumulation of zeaxanthin in B73. Additionally, a positive regulator of chilling stress, *CYTOKININ RESPONSE REGULATOR 1* (*CRR1*) (Zeng et al., 2021), was up-regulated in F7 and down regulated in B73 whereas the negative regulator of chilling stress, *MAP KINASE 8* (*MPK8*) (Zeng et al., 2021) showed the opposite expression (Fig. 2C and D). DESeq2 and TPM values for each of these genes can be found in Supp. dataset 1 and 2.

**Figure 2:**
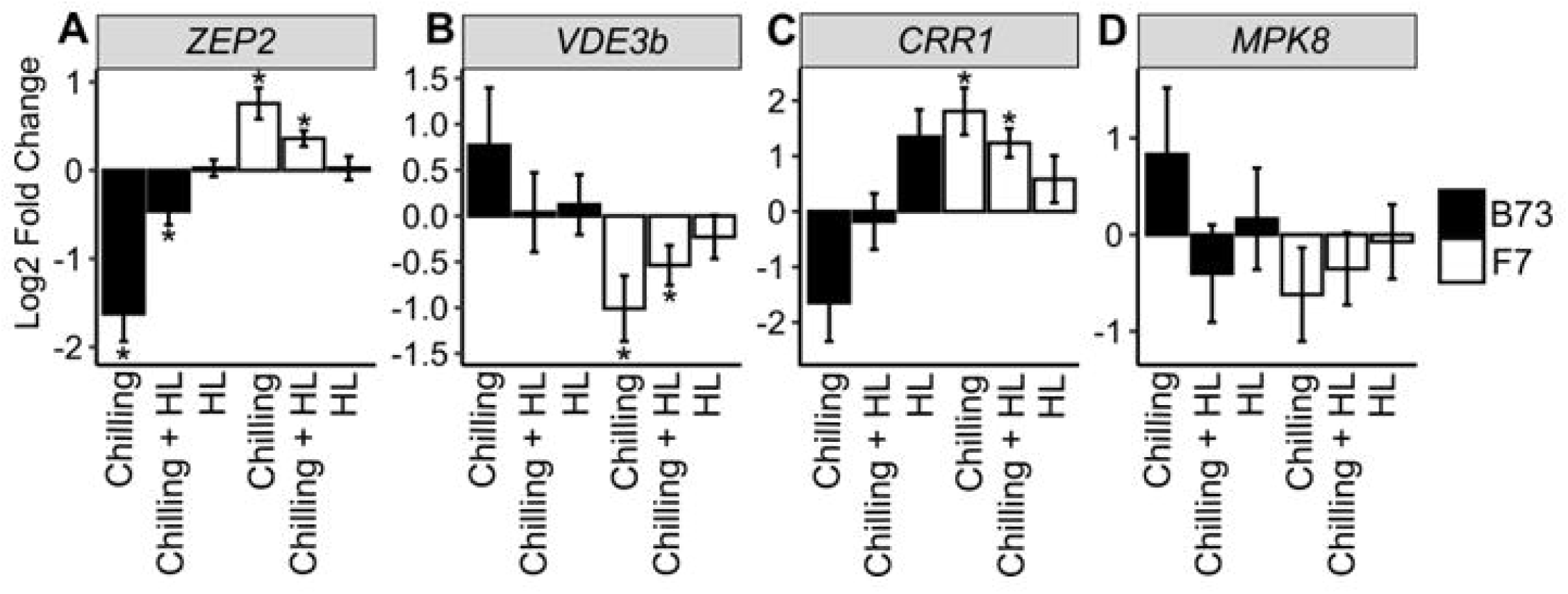
The expression of genes which indicate improved stress tolerance of F7 compared to B73. The Log2 Fold Change in expression of selected stress tolerance- related genes which show opposite patterns of expression in F7 compared to B73. Log2 Fold Change and *p-*adj values are from a DESeq2 analysis of the transcriptome data. Stars above bars indicate a statistically significant difference in Log2 Fold Change when compared with the untreated control (*p*-adj < 0.05 considered significant). Black and white bars indicate the B73 and F7 accessions respectively. DESeq2 and TPM values for each gene can be found in Supp. dataset 1 and 2. HL = High light. *ZEP2: ZEAXANTHIN EPOXIDASE 2* (Zm00001eb423570), *VDE3b: VIOLAXANTHIN DE-EPOXIDASE 3B* (Zm00001eb361500), *CRR1: CYTOKININ RESPONSE REGULATOR 1* (Zm00001eb066570), *MPK8: MAP KINASE 8* (Zm00001eb358890).

A recent study by Zhou et al. (2022a) carried out a transcriptome study on B73 and Mo17 accessions after a similar chilling treatment to that reported here. Mo17 is, like F7, considered more chilling tolerant than B73. To explore if mechanisms of chilling tolerance may be conserved between these two resilient accessions, genes differentially expressed in response to chilling in both F7 and Mo17, but not in B73, were determined. This resulted in 233 common genes, 133 up- regulated and 100 down-regulated (Supp. dataset 3). A GO enrichment analysis on these 233 genes showed “brassinosteroid metabolic process” as one of the most significantly enriched terms (FDR = 0.027), suggesting that a brassinosteroid-derived component of chilling tolerance (Ramirez and Poppenberger, 2020) may be common across these two accessions.

### High light mitigated the repressive effect of chilling on the transcription of photosynthetic genes

Rather than the hypothesized additive effect, the global transcriptome data suggested that high light had a mitigating effect on chilling responses (Fig. 1). To find groups of genes and subsequent individual candidate genes which may be involved in this response, a weighted gene co-expression network analysis (WGCNA) was performed. This analysis identified groups of genes (modules) which have similar patterns of expression. The eigengene values for the resulting 11 modules are shown in Supp. figure 2 (values provided in Supp. dataset 4) and provide a visual representation of the gene expression profiles in each module.

Modules one, two, three and four were of specific interest for further analyses The expression of the genes in modules one and two (Fig. 3A and B) showed large decreases in expression in response to the chilling treatment while this decrease in expression was not as severe (i.e. was mitigated) in the chilling plus high light treatment and there was minimal change in expression in response to high light alone. Thus, genes in these modules could be involved in the mitigating effect of high light on the severity of chilling stress. Based on the eigengene values, the expression of the genes in module one was similar across the two accessions (Fig. 3A) whilst genes in module two had slightly higher expression in B73 compared to F7 (Fig. 3B), although the pattern of expression was the same. To gain further biological insight into the role of the genes in these two modules, a GO enrichment analysis was performed on the gene lists from each module. This gave rise to multiple enriched GO terms in modules one and two (Supp. dataset 5). GO terms enriched in both modules were determined to identify possible physiological responses being most severely impacted by the chilling treatment and mitigated upon addition of high light (Supp. dataset 6). Several GO terms relating to photosynthesis were significantly enriched in modules one and two (Fig 3E.), including “photosystem II assembly”, ”chlorophyll metabolic process”, “thylakoid membrane organization”, and “photosynthetic electron transport chain”. Some other commonly enriched GO terms of interest here include “starch biosynthetic process”, “response to cold”, “membrane lipid biosynthetic process” and “response to oxygen-containing compound” (Fig. 3E). A heatmap of the Log2 Fold Change in expression of the genes with the “photosynthesis” GO terms found in modules one and two showed the progressive increase in expression from chilling to high light, with intermediate expression in the chilling plus high light treatment (Fig. 4A). Selected individual genes in Figure 4B exemplify the mitigating influence of high light on the chilling treatment, with all genes showing significant down-regulation in response to chilling and a notable increase in expression in the chilling plus high light treatment versus the chilling treatment. None of the genes showed a significant change in expression in response to high light alone. The pattern of expression of all genes is consistent between B73 and F7. These representative genes are involved in photosynthesis (*LCHB1, LHCB2*), NPQ (*PSBS*), carbon fixation (*PPDK1, PEPC1, RBCS*), xanthophyll metabolism (*VDE1, VP7*), chlorophyll metabolism (*CHAO2*), membrane integrity (*FAD6*), ROS scavenging (*SOD14*) and sucrose metabolism (*GAPA*) and give an example of the downregulated processes in the chilling treatment, for which high light addition seems to allow more sustained activity under chilling conditions. The DESeq2 and TPM values for each gene can be found in Supp. dataset 1 and 2. Overall, these results suggest that high light is able to mitigate the repressive effect of chilling on the transcription of genes primarily involved in photosynthesis and thus could improve growth in chilled environments.

**Figure 3:**
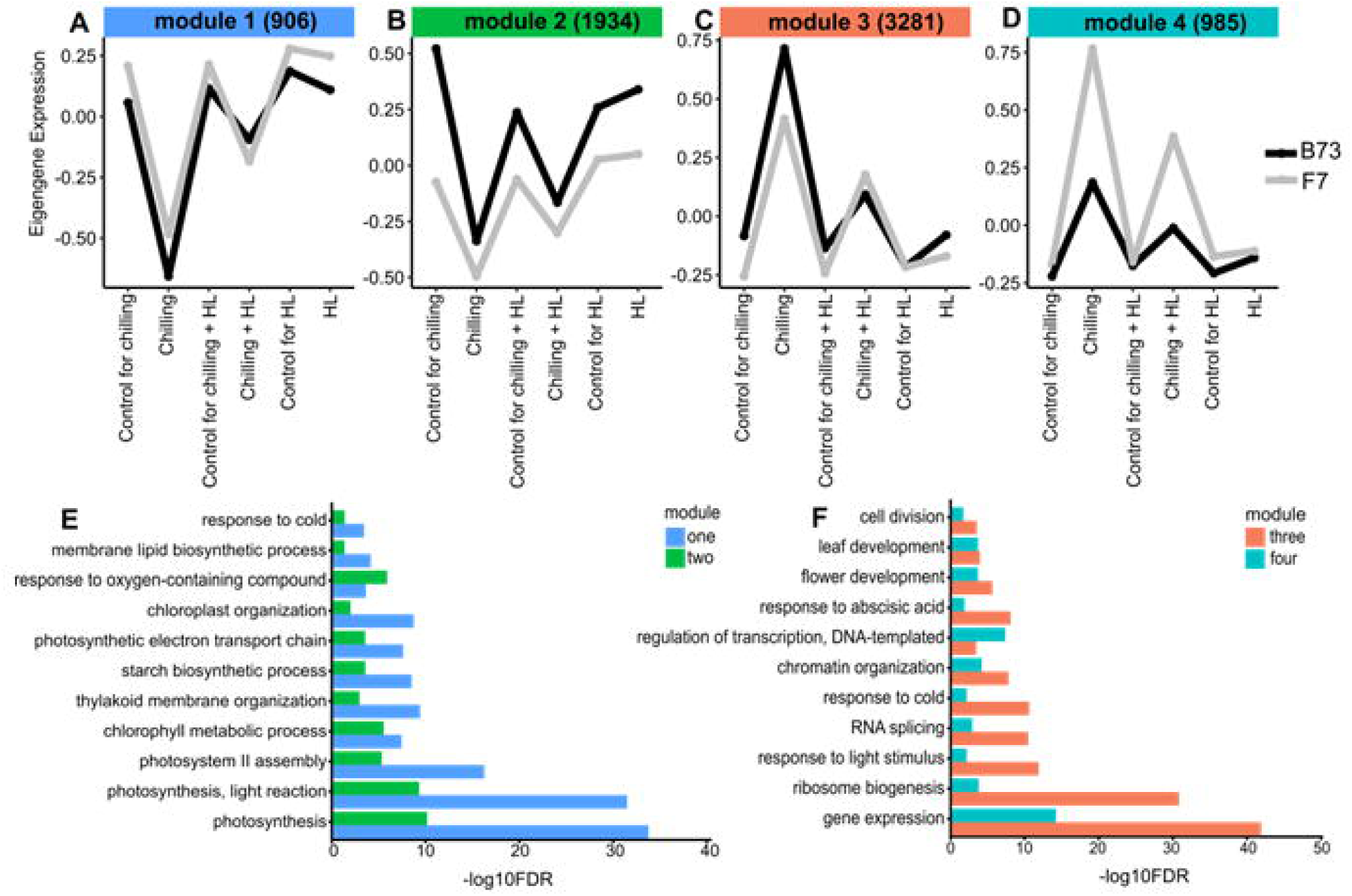
Eigengene expression of four modules from a WGCNA performed on the transcriptome data and GO enrichment analyses on the genes in the modules. **A, B, C** and **D** show the eigengene values representing the gene expression profiles of four out of the eleven modules identified by WGCNA. Numbers in brackets represent the number of genes in each of the modules. Lists of genes in each module can be found in Supp. dataset 4. **E**, GO terms of interest enriched in genes in module one and module two. **F**, GO terms of interest enriched in genes in module three and module four. For **E** and **F**, the significance of each GO term enrichment is represented as -log10FDR, with FDR representing false discovery rate.

**Figure 4:**
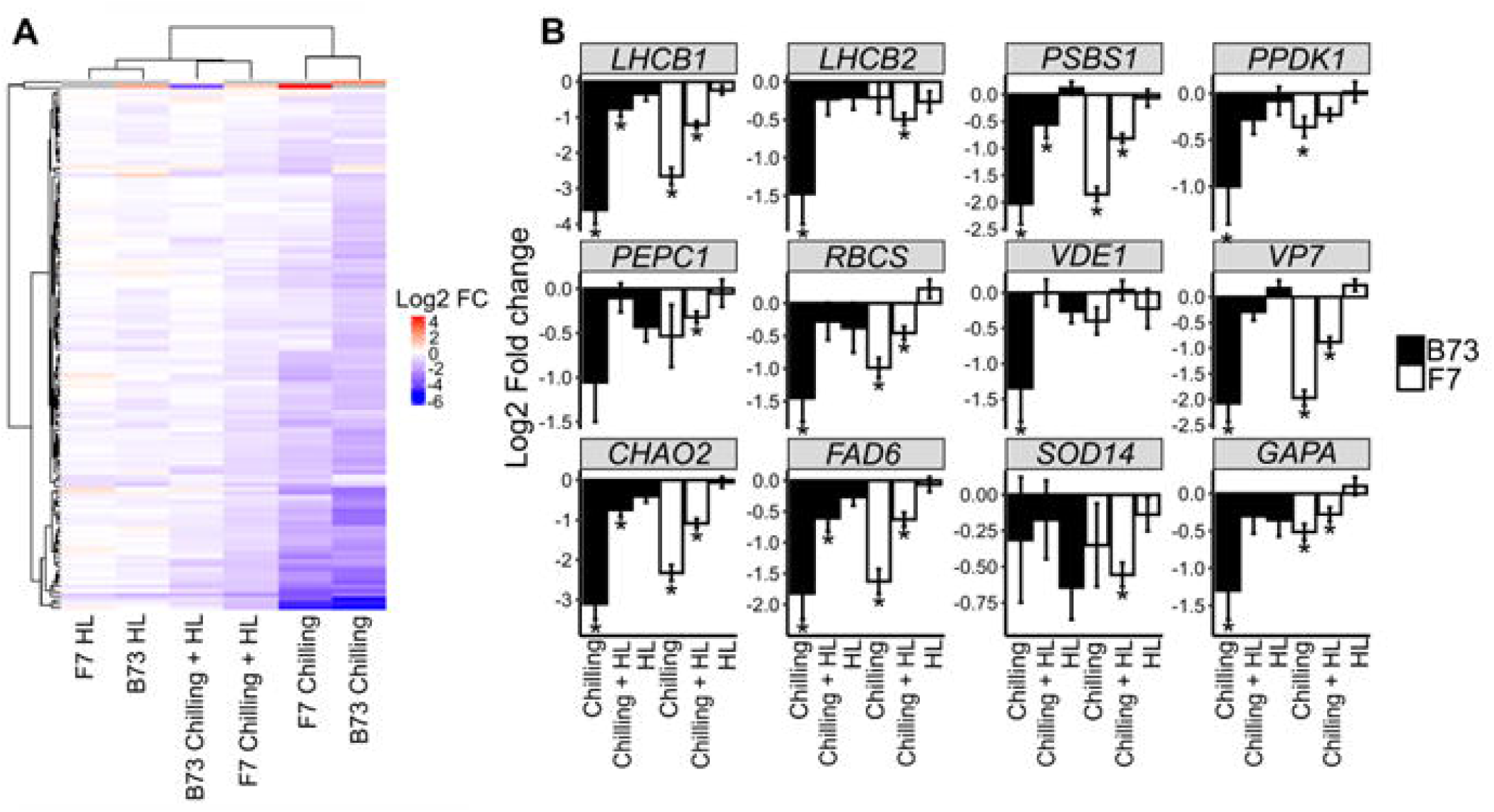
Heatmap and individual gene plots showing that the addition of high light to chilling treatment mitigates the repressive effect of chilling on gene expression in modules one and two. **A**, a heatmap of the Log2 Fold Change (FC) in expression of genes found in modules one and two with the “Photosynthesis” GO term (82 genes from module one and 75 genes from module two). **B**, the Log2 Fold Change in expression of selected genes found in modules one or two involved in photosynthesis (*LCHB1, LHCB2*), NPQ (*PSBS*), carbon fixation (*PPDK1, PEPC1, RBCS*), xanthophyll metabolism (*VDE1, VP7*), chlorophyll metabolism (*CHAO2*), membrane integrity (*FAD6*), ROS scavenging (*SOD14*) and sucrose metabolism (*GAPA*). Log2 Fold Change and *p*-adj values are from a DESeq2 analysis of the transcriptome data. Stars above bars indicate a statistically significant difference in Log2 Fold Change when compared with the untreated control (*p*-adj < 0.05 considered significant). Black and white bars indicate the B73 and F7 accessions respectively. DESeq2 and TPM values for each gene can be found in Supp. Dataset 1 and 2. *LCHB1: LIGHT HARVESTING CHLOROPHYLL A/B BINDING PROTEIN 1* (Zm00001eb296090)*, LHCB2: LIGHT HARVESTING CHLOROPHYLL A/B BINDING PROTEIN 2* (Zm00001eb320890)*, PSBS: PHOTOSYSTEM II SUBUNIT* (Zm00001eb146510)*, PPDK1: PYRUVATE, PHOSPHATE DIKINASE 1* (Zm00001eb287770)*, PEPC1: PHOSPHOENOLPYRUVATE CARBOXYLASE 1* (Zm00001eb383680)*, RBCS: RUBISCO SMALL SUBUNIT* (Zm00001eb092540)*, VDE1: VIOLAXANTHIN DE-EPOXIDASE 1* (Zm00001eb085840)*, VP7: VIVIPAROUS 7* (Zm00001eb234930)*, CHAO2: CHLOROPHYLLIDE A OXYGENASE 2* (Zm00001eb362510)*, FAD6: FATTY ACID DESATURASE 6* (Zm00001eb041150)*, SOD14: SUPEROXIDE DISMUTASE 14* (Zm00001eb039000)*, GAPA: GLYCERALDEHYDE 3- PHOSPHATE DEHYDROGENASE SUBUNIT A* (Zm00001eb080840).

### High light mitigated the inducing effect of chilling on the transcription of ABA signalling, plant growth and transcription related genes

As with modules one and two, modules three and four (Fig. 3C and D) were of specific interest for further analysis as they also showed a mitigating effect of high light when combined with a chilling treatment, but in the opposite direction to modules one and two. The expression of the genes in these modules showed large increases in expression in response to the chilling treatment but this was weaker in the chilling plus high light treatment and there was minimal change in expression in response to high light alone. Based on eigengene values, expression of genes in module three was similar across the two accessions whilst genes in module four had higher expression in F7 compared to B73, although the pattern of expression was the same. GO enrichment analysis of the module gene lists showed that the “gene expression” GO term as one of the most significantly enriched term in both gene lists (Fig. 3F, Supp. dataset 5) several GO terms related to gene expression, including “regulation of transcription, DNA templated”, “RNA splicing”, “ribosome biogenesis” and “chromatin organization”. Other GO terms of interest enriched in both module gene lists included “response to abscisic acid”, “leaf development” and “flower development”. Figure 5A shows a heatmap of the Log2 Fold Change in expression of the genes with the “regulation of transcription, DNA-templated” GO term. Expression of these genes is most up-regulated in the chilling treated samples, whilst this up-regulation is decreased in the chilling plus high light samples. As with modules one and two above (Fig. 4B), several genes representative of transcript abundance and overall enriched processes in modules three and four were graphed for clearer visualisation (Fig. 5B). Each gene was significantly up-regulated in response to chilling and a notable decrease in the expression in response to chilling plus high light was evident. Again, there were no statistically significant changes in expression in response to high light alone. Genes in figure 5B are involved in processes including ABA signalling (*SnRK2.9, CDPK4, CDPK7, PYL8, PYL13*), stress responsive transcription (*DREB1.2, DREB1.7*), regulation of translation (*AGO1A, AGO1B*) and leaf senescence (*S40-1, S40-2, S40-4*). The DESeq2 and TPM values for each gene can be found in Supp. dataset 1 and 2. These results suggest that the addition of high light during chilling dampens the chilling-enhanced transcription of genes modulating ABA signalling, growth and transcription and which could improve growth in chilled environments.

**Figure 5:**
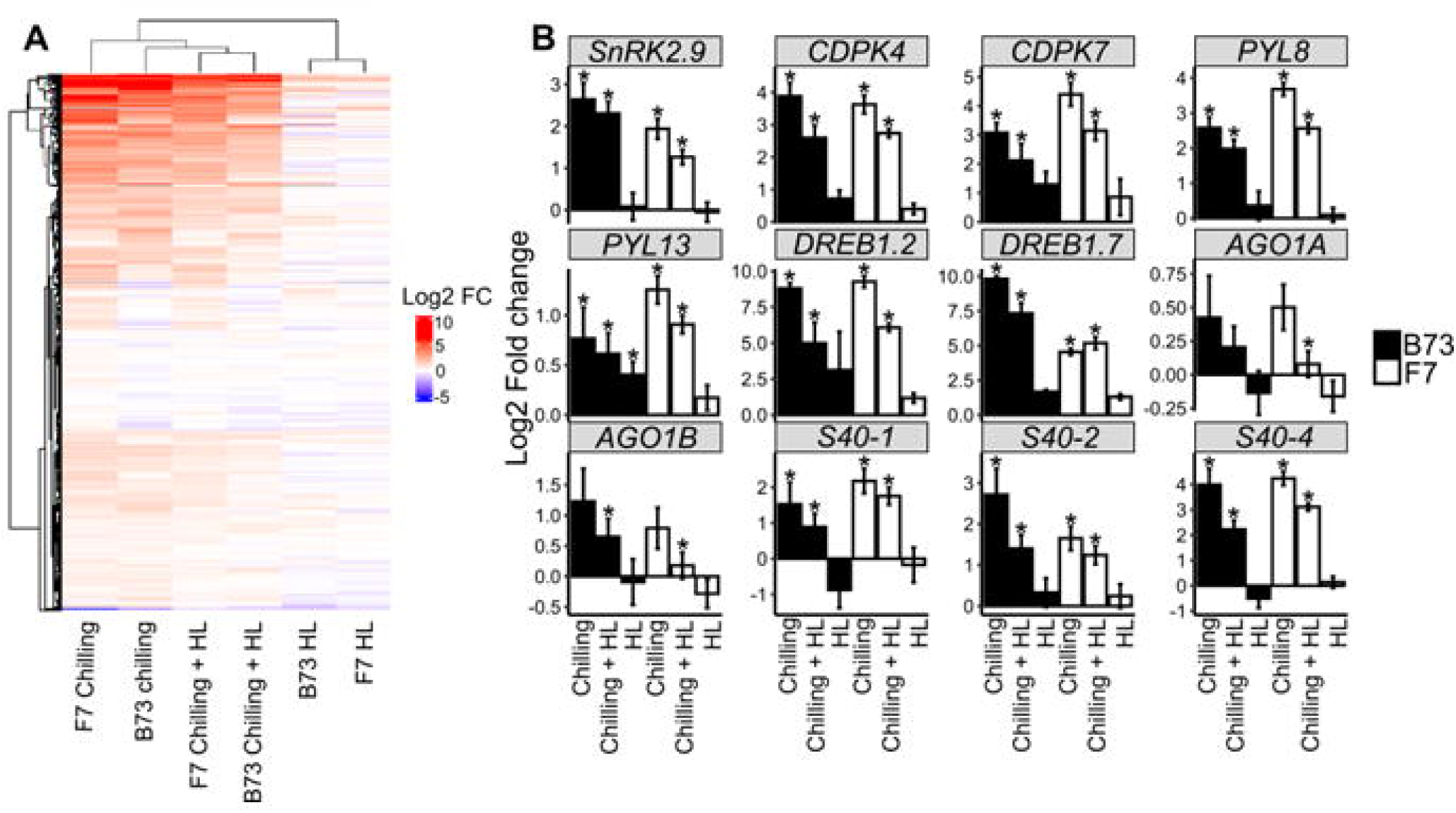
Heatmap and individual gene plots showing that the addition of high light to chilling treatment mitigates the inducing effect of chilling on gene expression in modules three and four. **A**, a heatmap of the Log2 Fold Change (FC) in expression of genes in modules three and four with the “regulation of transcription, DNA-templated” GO term (604 genes from module three and 228 genes from module four). **B**, the Log2 Fold Change in expression of selected genes found in modules three or four involved in ABA signalling (*SnRK2.9, CDPK4, CDPK7, PYL8, PYL13*), stress responsive transcription (*DREB1.2, DREB1.7*), regulation of translation (*AGO1A, AGO1B*) and leaf senescence (*S40-1, S40-2, S40-4*). Log2 Fold Change and *p*-adj values are from a DESeq2 analysis of the transcriptome data. Stars above bars indicate a statistically significant difference in Log2 Fold Change when compared with the untreated control (*p*-adj < 0.05 considered significant). Black and white bars indicate the B73 and F7 accessions respectively. DESeq2 and TPM values for each gene can be found in Supp. dataset 1 and 2. *SNRK2.9: SNF1- RELATED PROTEIN KINASE 2.9* (Zm00001eb051510), *CDPK4: CALCIUM DEPENDENT PROTEIN KINASE 4* (Zm00001eb428080), *CDPK7: CALCIUM DEPENDENT PROTEIN KINASE 7* (Zm00001eb187630), *PYL8: PYRABACTIN RESISTANCE-LIKE PROTEIN 8* (Zm00001eb350630), *PYL13: PYRABACTIN RESISTANCE-LIKE PROTEIN 13* (Zm00001eb204180), *DREB1.2: DEHYDRATION-RESPONSIVE ELEMENT-BINDING PROTEIN 1.2* (Zm00001eb318890), *DREB1.7: DEHYDRATION-RESPONSIVE ELEMENT- BINDING PROTEIN 1.7* (Zm00001eb269420*), AGO1A: ARGONAUTE 1A* (Zm00001eb267430), *AGO1B: ARGONAUTE LINKER 2 DOMAIN* (Zm00001eb428910), *S40-1: SENESCENCE REGULATOR S40-1* (Zm00001eb077160), *S40-2: SENESCENCE REGULATOR S40-2* (Zm00001eb083770), *S40-4: SENESCENCE REGULATOR S40-4* (Zm00001eb221600).

### Identification of candidate transcription factors driving the high light mitigation of chilling stress

Transcription factors (TFs) play a vital role in the response to all abiotic stresses, including chilling and high light (reviewed by Abdullah, Azzeme and Yousefi, 2022, Sharma et al., 2020, Wang et al., 2016). With this in mind, the TFs and enriched TF binding motifs present in the WGCNA module gene lists were investigated to find candidates that could be responsible for the mitigating effect of high light treatment on the transcriptome signature and stress response of chilling.

Firstly, the TFs present in each module gene list were identified using the iTAK tool (Zheng et al., 2016). A total of 39 TFs were identified in the gene list from module one and 88 TFs identified in module two (Table 1). In both modules, the most represented types of TF were homeobox (HB) TFs (9 and 8 present in modules one and two respectively) and MYB-related TFs (4 and 10 present in modules one and two respectively). Both modules also contained TFs from bZIP, C3H and GATA TF families. Only module two contained TFs from the heat shock factor (HSF) family. A total of 213 TFs were identified in the gene list from module three and 78 TFs identified in module four (Table 2). APETALA2/Ethylene Responsive Element Binding Factor domain protein (AP2/EREB) TFs were the most represented type of TF in both module three and four (24 and 9 present in module three and four respectively). Other TFs identified in both modules included WRKY, bZIP and NAC TF families. As expected, most of the highly represented TF families from modules three and four were different from those in modules one and two. All of the highly represented TF families in both gene lists are well known for modulating abiotic stress responses (Abdullah, Azzeme and Yousefi, 2022, Sharma et al., 2020, Wang et al., 2016).

**Table 1:** Transcription factors present in modules one and two. The No. column refers to the number of transcription factors of that type present in the list. For a comprehensive list of gene IDs and “Other TFs”, refer to Supp. dataset 7.

| Type of TF | No. | Description |
| --- | --- | --- |
| **Module one** |  |  |
| HB | 9 | Homeobox, homeodomain, leucine-zipper containing protein |
| MYB-related | 4 | MYB domain protein |
| C3H | 4 | Zinc finger (CCCH-type) family protein |
| GRAS | 3 | GRAS family transcription factor |
| NF | 3 | Nuclear transcription factor (NF-X1, NF-YA, NF-YB, NF-YC) |
| C2C2-GATA | 3 | Class IV zinc finger transcription factor |
| bZIP | 1 | Basic -leucine zipper protein |
| Other TFs | 12 |  |
| **Total TFs** | **39** |  |
| **Module two** |  |  |
| MYB-related | 10 | MYB domain protein |
| HB | 8 | Homeobox, homeodomain, leucine-zipper containing protein |
| bZIP | 8 | Basic -leucine zipper protein |
| C3H | 7 | Zinc finger (CCCH-type) family protein |
| C2H2 | 7 | C2H2-type zinc finger family protein |
| GeBP | 5 | Leucine-zipper transcription factors |
| HSF | 5 | Heat shock factor |
| NF | 3 | Nuclear transcription factor (NF-X1, NF-YA, NF-YB, NF-YC) |
| GRAS | 2 | GRAS family transcription factor |
| C2C2-GATA | 2 | Class IV zinc finger transcription factor |
| Other TFs | 41 |  |
| **Total TFs** | **88** |  |

**Table 2:** Transcription factors present in modules three and four. The No. column refers to the number of transcription factors of that type present in the list. For a comprehensive list of gene IDs and “Other TFs”, refer to Supp. dataset 7.

| Type of TF | No. | Description |
| --- | --- | --- |
| **Module three** |  |  |
| AP2/EREB | 24 | APETALA2/Ethylene Responsive Element Binding Factor domain protein |
| WRKY | 19 | WRKY family transcription factor |
| NAC | 18 | NAM, ATAF and CUC (NAC) domain-containing protein |
| C3H | 15 | Zinc finger (CCCH-type) family protein |
| GRAS | 13 | GRAS family transcription factor |
| C2H2 | 12 | C2H2-type zinc finger family protein |
| C2C2-Dof | 10 | Zinc finger transcription factor, DNA-binding with one finger |
| MYB-related | 8 | MYB domain protein |
| Trihelix | 8 | Trihelix transcription factor |
| bZIP | 5 | Basic -leucine zipper protein |
| Other TFs | 81 |  |
| **Total TFs** | **213** |  |
| **Module four** |  |  |
| AP2/EREB | 9 | APETALA2/Ethylene Responsive Element Binding Factor domain protein |
| MYB-related | 9 | MYB domain protein |
| C2C2-Dof | 4 | Zinc finger transcription factor, DNA-binding with one finger |
| bZIP | 4 | Basic -leucine zipper protein |
| NAC | 3 | NAM, ATAF and CUC (NAC) domain-containing protein |
| C3H | 3 | Zinc finger (CCCH-type) family protein |
| GRAS | 2 | GRAS family transcription factor |
| Trihelix | 2 | Trihelix transcription factor |
| WRKY | 1 | WRKY family transcription factor |
| Other TFs | 41 |  |
| **Total TFs** | **78** |  |

Following the identification of TFs in each module, a Simple Enrichment Analysis (SEA) using the MEME Suite of Motif-based sequence analysis tools was performed to identify enriched TF binding motifs present in the promoters of genes in each module, using 1000 bp of genomic sequence located upstream of the ATG start codon of each gene. Motif enrichment was assessed in comparison to a control set of 5036 genes which did not show significant differential expression in any of the treatments. The promoters of the genes in modules one and two had 28 and 40 enriched TF binding motifs respectively (Table 3). The majority of these were from the bHLH TF family (10 in module one and 17 in module two) and the bZIP TF family (10 in module one and 13 in module 2). Motifs for the HSF TF family were also enriched in modules one and two (1 on module one and 6 in module two). The promoters of the genes in modules three and four had 52 and 18 enriched TF binding motifs respectively (Table 4). Unlike modules one and two, the types of TF families with enriched promoter binding sites in modules three and four differed, with the AP2/EREB TF family being most represented in module three (24 enriched motifs) and the bHLH TF family being most represented in module four (7 enriched motifs).

**Table 3:** The transcription factor binding motifs enriched in the promoters of genes in modules one and two. The No. column refers to the number of genes with that specific TF binding sight enriched. For CIS-BP IDs, gene IDs and enrichment data refer to Supp. dataset 8.

| TF Family name | No. | Description |
| --- | --- | --- |
| **Module one** |  |  |
| bHLH | 10 | Basic helix-loop-helix |
| bZIP | 10 | Basic -leucine zipper protein |
| SBP | 3 | SQUAMOSA promoter-binding protein |
| HSF | 1 | Heat shock factor |
| Myb/SANT | 1 | MYB domain protein |
| Other | 3 |  |
| **Total enriched TF motifs** | **28** |  |
| **Module two** |  |  |
| bHLH | 17 | Basic helix-loop-helix |
| bZIP | 13 | Basic -leucine zipper protein |
| HSF | 6 | Heat shock factor |
| NAC | 1 | NAM, ATAF and CUC transcription factors |
| Other | 3 |  |
| **Total enriched TF motifs** | **40** |  |

**Table 4:**
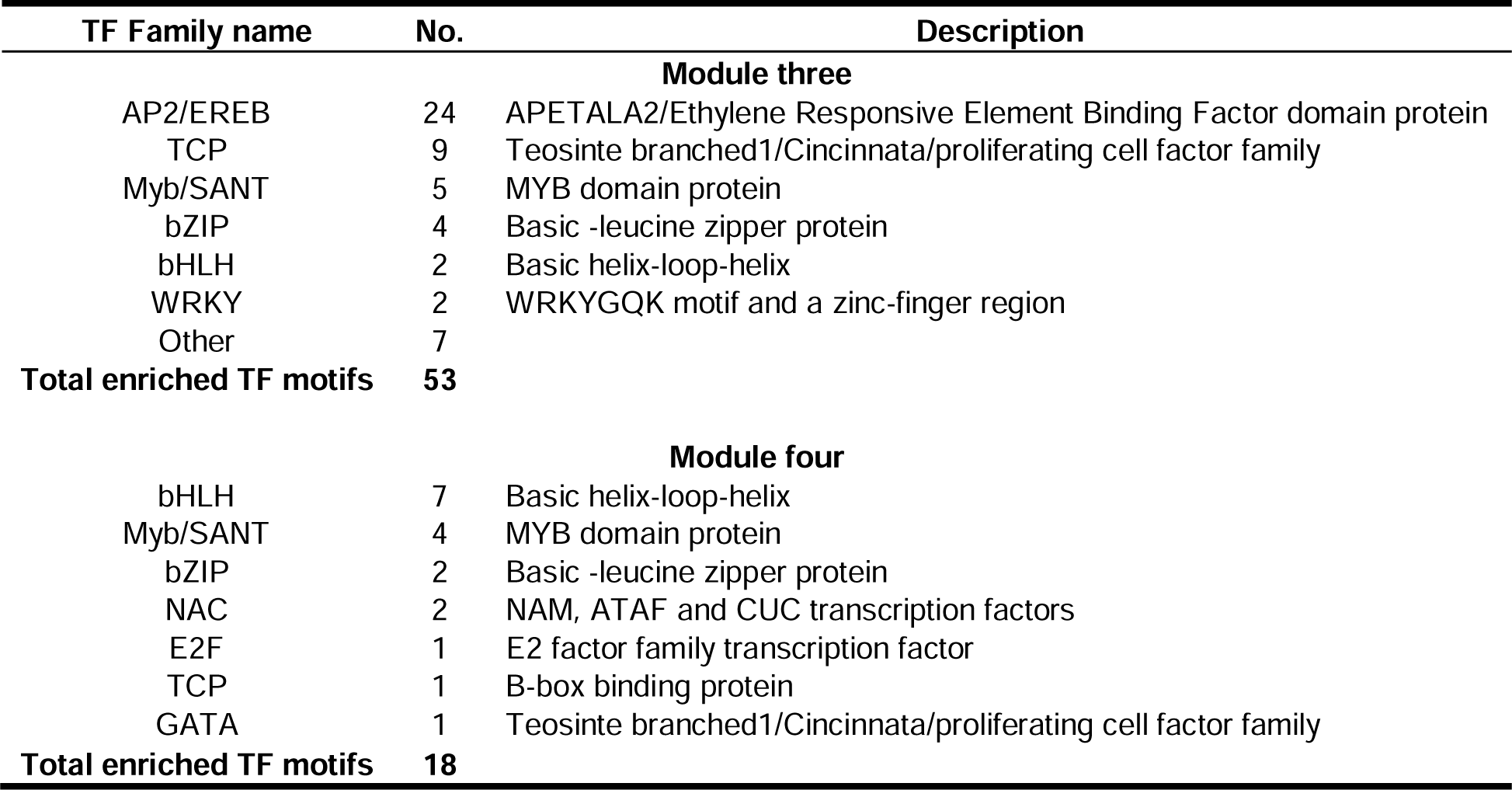
The transcription factor binding motifs enriched in the promoters of genes in modules three and four. The No. column refers to the number of genes with that specific TF binding sight enriched. For CIS-BP IDs, gene IDs and enrichment data refer to Supp. dataset 8.

**Table 5:** Information on the candidate transcription factors selected based on their presence in modules one, two, three or four and an enriched transcription factor binding motif in the promoters of the other genes in the module. The references column includes work illustrating the role of each TF, or the TF family in which they belong,

| GeneID | Gene name | Functional annotation | Module | kME for module | Rank in SEA | Previous studies |
| --- | --- | --- | --- | --- | --- | --- |
| Zm00001eb339360 | bHLH99 | Basic helix-loop-helix transcription factor | 2 | 0.95 | 19/40 | - Li et al ., 2019<br>- Li et al ., 2021 |
| Zm00001eb301280 | HSF4 | Heat stress transcription factor 4 | 2 | 0.91 | 1/40 | - Sowinski et al ., 2020<br>- Zhou et al ., 2021<br>- Lui et al ., 2022 |
| Zm00001eb037480 | TCP21 | TCP-transcription factor 21 | 3 | 0.95 | 14/53 | - Ding et al , 2019<br>- Danisman, 2016 |
| Zm00001eb364110 | TCP20 | TCP-transcription factor 20 | 3 | 0.91 | 17/53 | - Ding et al , 2019<br>- Danisman, 2016 |
| Zm00001eb145420 | EREB52 | AP2-EREBP-transcription factor 52 | 3 | 0.95 | 4/53 | - Ritonga et al ., 2021<br>- Zhou et al ., 2022<br>- Qin et al ., 2004<br>- Wang & Dong, 2009<br>- Wang, Yang & Yang, 2011 |

during chilling and/or high light responses

To identify candidate transcription factors that may underpin the mitigation of chilling stress by high light, the results from the iTAK analysis (Tables 1 and 2) and SEA (Tables 3 and 4) were combined. Here, the gene IDs of the top 20 enriched TF binding motifs from the Simple Enrichment Analysis (SEA) results were scanned (except for module four where only 18 were identified) to determine whether they were present in the list of TFs in the iTAK output (i.e. a transcription factor present in module one which also had an enriched binding motif in the promoters of the other genes in module one). The subsets of transcription factors were then filtered for a module membership (kME) value greater than 0.9. The kME values indicate the centrality of a gene within a WGCNA module, with the most central ‘hub’ genes within a module having a kME value approaching 1. This resulted in the identification of five TFs, present only in module two or three (Table 5 and Figure 6). Heatmaps in Figure 6 represent the 18-20 most enriched TF binding motifs in the four modules of interest, the TF family of each motif indicated by the text to the right of the corresponding heatmap box. The five TF candidates identified are highlighted by black boxes within the heatmap and the DESeq2 expression results included adjacent. *HEAT STRESS TRANSCRIPTION FACTOR 4 (HSF4)* and (*BASIC HELIX-LOOP-HELIX TRANSCRIPTION FACTOR 99 (BHLH99)* were identified from module two. And *AP2-EREBP-TRANSCRIPTION FACTOR 52 (EREB52), TEOSINTE- BRANCHED1/CYCLOIDEA/PCF TRANSCRIPTION FACTOR 21 (TCP21)* and *TEOSINTE-BRANCHED1/CYCLOIDEA/PCF TRANSCRIPTION FACTOR 20 (TCP20)* were identified from module three. Assuming that expression changes of these TF encoding genes are indicative of expression changes of the target genes which they regulate (i.e. the genes present in each module), the five TFs identified above serve as candidate transcription factors that modulate the mitigating effect which high light has on chilling treatment and may have potential to improve maize growth in chilled conditions.

**Figure 6:**
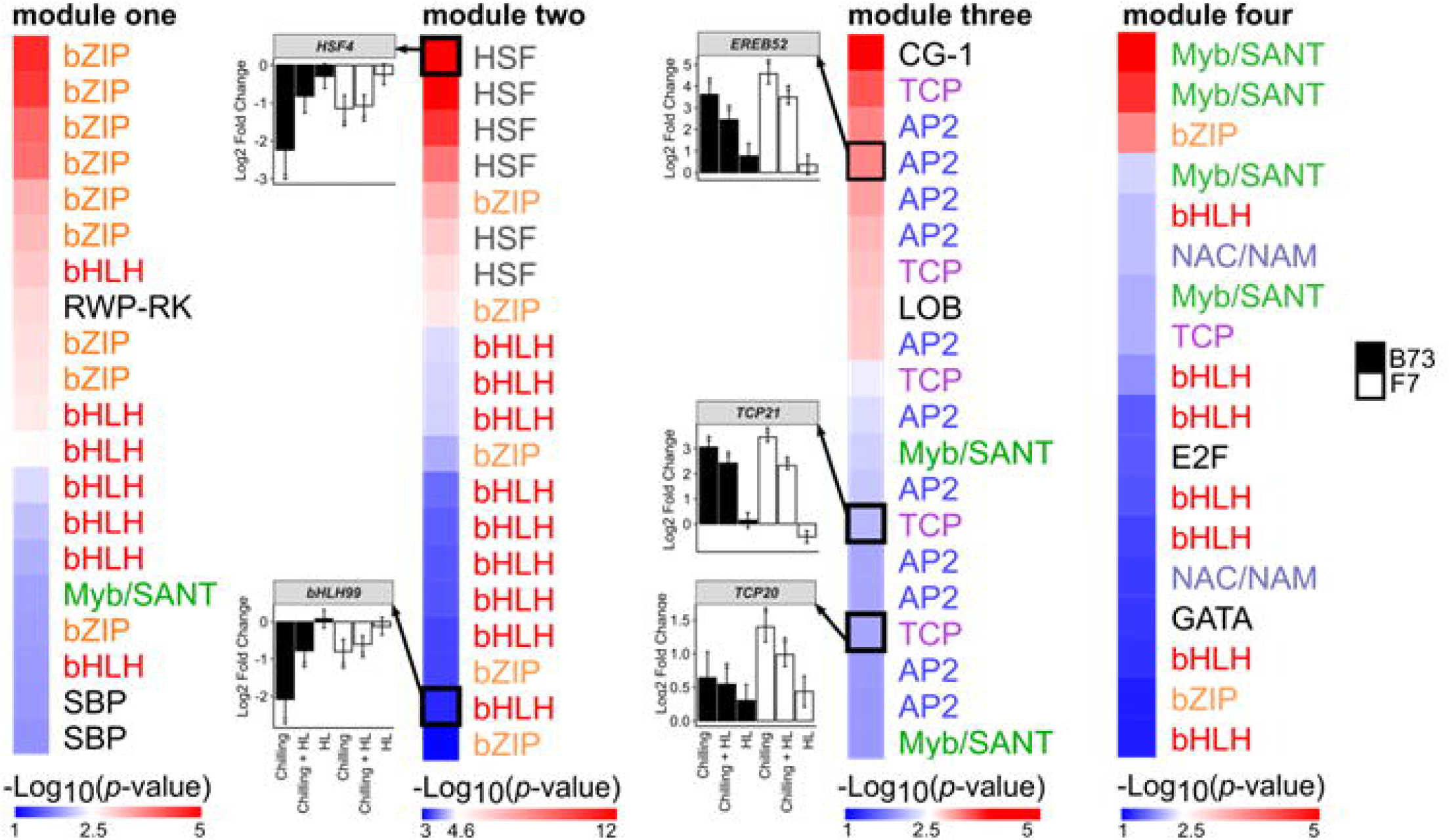
Identification of candidate transcription factors driving the mitigating effect of high light on the transcriptomic response to chilling stress. The heatmaps show the 20 most enriched transcription factor binding motifs present in the promoters of genes in modules one, two, three and four. Enriched motifs were identified using a Simple Enrichment Analysis (SEA) on sequences 1000 bp upstream of the ATG transcription start site. The TF family of each TF binding motif is indicated next to the corresponding box in each heatmap. Font colour groups TFs according to TF family. The degree of motif enrichment is depicted through -Log_10_(*p-value*). The transcription factors highlighted with a black box indicate those which had enriched binding motifs identified in the SEA, were also present in the genes list for the corresponding module and had a module kME greater than 0.9. Bar plots show the expression of the identified TF candidates from a DESeq2 analyses of the transcriptome data. Stars above bars indicate a statistically significant difference in Log2 Fold Change when compared with the untreated control (*p*-adj < 0.05 considered significant). Black and white bars indicate the B73 and F7 accessions respectively. DESeq2 and TPM values for each gene can be found in Supp. dataset 1 and 2. *HSF4*: *HEAT STRESS TRANSCRIPTION FACTOR 4* (Zm00001eb301280)*, BHLH99*: *BASIC HELIX-LOOP-HELIX TRANSCRIPTION FACTOR 99* (Zm00001eb339360)*, EREB52: AP2-EREBP-TRANSCRIPTION FACTOR 52* (Zm00001eb145420)*, TCP21: TEOSINTE-BRANCHED1/CYCLOIDEA/PCF TRANSCRIPTION FACTOR 21* (Zm00001eb037480), *TCP20: TEOSINTE- BRANCHED1/CYCLOIDEA/PCF TRANSCRIPTION FACTOR 20* (Zm00001eb364110).

### Decrease in F_v_/F_m_ due to chilling stress was alleviated by high light

A number of common physiological parameters were measured in response to each treatment to find out if these would be consistent with the aforementioned transcriptomics-based observations. No detectable effect of chilling, chilling plus high light or high light alone was found on the concentration of malondialdehyde (MDA) which is a measure of the lipid peroxidation state (Supplemental Fig. 3), or on the chlorophyll a/b ratio (Supplemental Fig. 4). There was an increase in total chlorophyll content compared to the control under the high light treatment but there was no effect of chilling or chilling plus high light on chlorophyll content (Supplemental Fig. 4). Chlorophyll fluorescence imaging was used to assess the impact of chilling and high light treatments on the photosynthetic apparatus. Leaf discs taken from plants exposed to chilling and/or high light stress were dark adapted for 1 hour after which F_v_/F_m_ was measured; this provided an estimate of the maximum quantum efficiency of photosystem II. F_v_/F_m_ was significantly decreased by 9-12% in the chilling- exposed plants (*p*-value < 0.001, Fig. 7A) indicating damage to photosystem II. F_v_/F_m_ decreased much less in leaves exposed to chilling stress in combination with high light (2-3% decrease, *p*-value < 0.05, Fig. 7B) and in leaves exposed to high light alone (3% decrease, *p*-value < 0.05, Fig. 7C).

**Figure 7:**
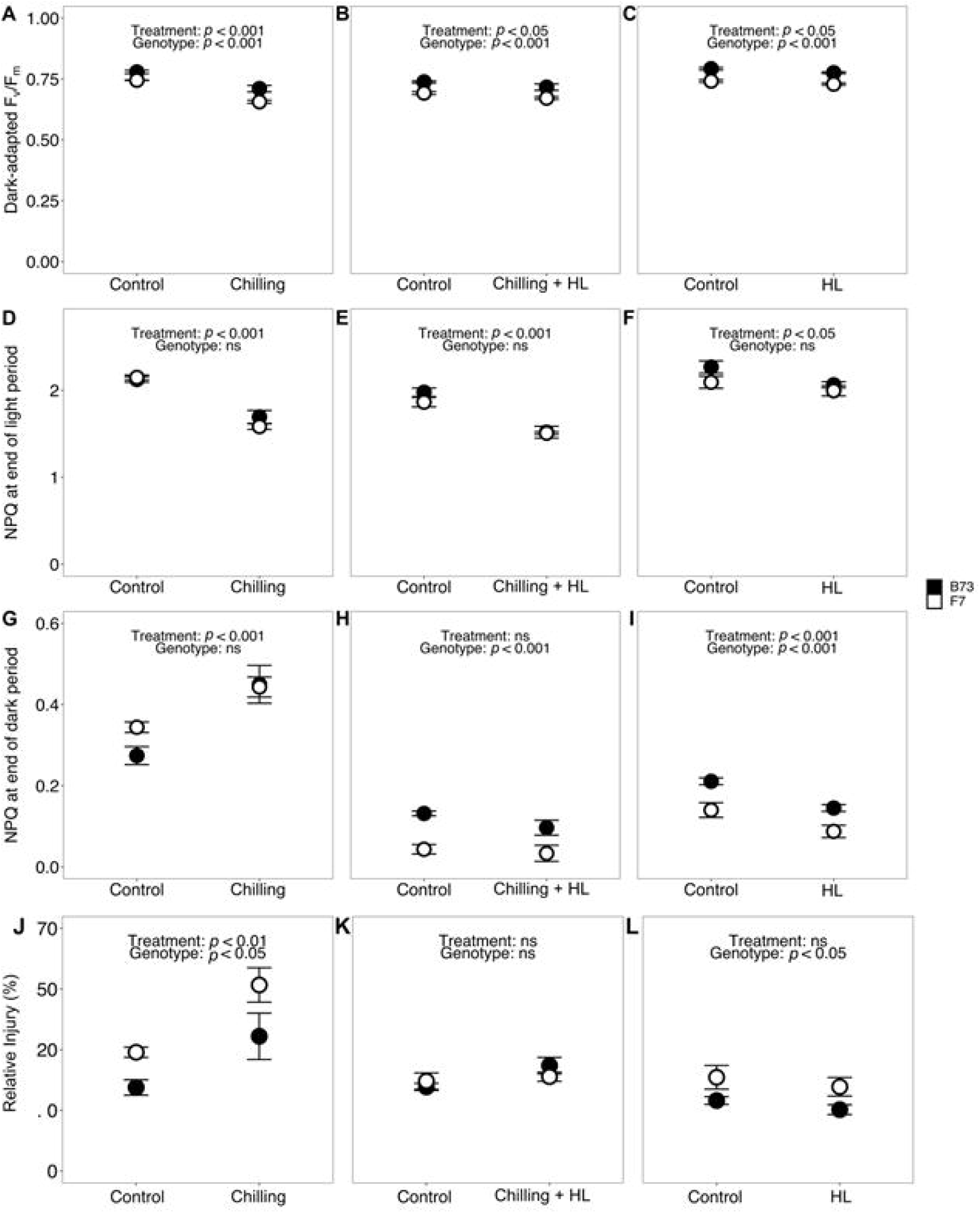
High light mitigates changes in F_v_/F_m_, NPQ and relative injury resulting from chilling stress. **A-I**, leaf discs sampled from maize plants that had previously been given chilling, chilling plus high light (HL), high light (HL) treatments, or corresponding control treatments, were exposed to 10 minutes of light and 12 minutes of dark recovery. **A**, **B**, **C**, Dark-adapted F_v_/F_m_; **D**, **E**, **F**, NPQ level at the end of 10 min light; **G**, **H**, **I**, NPQ level at the end of 12 min dark recovery. **J**, **K**, **L**, relative injury of leaf discs sampled from maize plants that had previously been given chilling, chilling plus high light, high light treatments, or corresponding control treatments. Each treatment was analysed with respect to its own control (n=4). Results from two-way ANOVA are shown. There were no statistically significant interactions between accession and treatment.

### Chilling stress impaired photoprotective responses

Following measurement of F_v_/F_m_, leaf discs were immediately exposed to a 10 minute light treatment and a 12 minute dark relaxation period to assess how the ability to dissipate absorbed energy via photochemical and nonphotochemical quenching processes was affected by the treatments. Non-photochemical quenching (NPQ) is the photoprotective downregulation of light-harvesting efficiency, allowing dissipation of excess light energy not used in photochemistry. Significantly less NPQ induction occurred in leaves that had been exposed to the chilling treatment compared to the relevant control: amplitude of NPQ at the end of the light phase was 20-26% lower (*p*-value < 0.001, Fig. 7D). This was also the case for chilling stress in combination with high light (19-23% decrease, *p*-value < 0.001, Fig. 7E). There was a marginal decrease in NPQ at the end of the light phase (5-9%, *p*-value < 0.05, Fig. 7F) in leaves that had been subjected to the high light treatment compared to the relevant control. During the 12 minute dark relaxation phase, significantly less NPQ relaxation occurred in the leaves that had been exposed to the chilling treatment, leading to higher residual NPQ at the end of the dark phase (29-64% increase, *p*- value < 0.001, Fig. 7G). However, in leaves that had been exposed to combined chilling and high light, there was no significant difference in residual NPQ between treatment and control (Fig. 7H) suggesting that the addition of high light allowed a more complete relaxation of NPQ in the 12 minute dark relaxation. In leaves that had been exposed to high light only, residual NPQ was 31-37% lower than in the control (*p*-value < 0.001, Fig. 7I). Again, these results confirm the findings in the transcriptome data indicating that high light is able to mitigate chilling stress.

### Relative injury was increased by chilling stress but mitigated by high light

In plants exposed to chilling, the relative injury of leaf tissue, a measure of stress-induced electrolyte leakage due to membrane breakdown, was 57-61% greater than in control plants (*p*-value < 0.01, Fig. 7J). When plants were exposed to chilling stress in combination with high light, relative injury was not different than the control (Fig. 7K) suggesting that high light mitigated the chilling-induced damage, in line with the transcriptome data above. Relative injury in response to high light without chilling also did not differ from the control (Fig. 7L).

## Discussion

Contrasting results have been reported on the effect of adding high light to chilling-induced stress in cold-sensitive plants like maize. Some work suggests that high light would add to chilling-induced stress, primarily via photoinhibition and photodamage (Farage and Long, 1987, Ortiz-Lopez, 1990), whereas others suggest that high light can aid acclimation to chilling stress and thereby mitigate the negative effects of chilling (Szalai et al., 2018, Grzybowski et al., 2019). These discrepancies could be due to different chilling and high light treatments and growth environments, thus making direct comparisons of the results difficult. The results presented here show that the addition of three hours of high light in the morning as part of a 12 h night and 3 hr morning chilling treatment had a surprisingly consistent mitigating impact on the effect of chilling on both transcriptomic profiles and physiological status of the leaves (Fig. 1, 3, 4, 5, and 7). These responses were found in both B73 and F7 accessions, although the transcriptomic responses of F7 indicated that it is more tolerant to chilling than B73, consistent with previous reports (Enders et al., 2018).

### The role of high light in mitigating chilling stress

The transcriptome and physiological data suggest that three hours of increased light in the morning had very little impact as an individual treatment but was significantly more impactful when plants had already had 12 hours of cold, dark treatment. Modules one, two, three and four of the WGCNA contained genes with patterns of expression which depicted the mitigating effect of high light on the chilling treatment (Fig. 3A, B, C and D). GO enrichment analyses and analyses of individual genes in modules one and two showed that processes such as photosynthesis, xanthophyll metabolism, chlorophyll metabolism, response to ROS and sucrose responses were all decreased in response to chilling but mitigated by high light treatment (Fig. 3E, Fig. 4A). These physiological responses have previously been reported as repressed in chilling environments and result in damaging physiological changes and reduced plant growth (reviewed by Burnett and Kromdijk, 2022, Savitch et al., 2011). Based on the expression patterns of many genes with these GO terms, it is plausible that the addition of high light was able to reduce the severity of chilling-induced down-regulation of these processes, thus improving physiological integrity. F_v_/F_m_ and NPQ relaxation results (Fig. 7) substantiate that the addition of high light alleviates chilling-induced strain on photosynthesis and photoinhibition. GO enrichment analyses and analyses of individual genes in modules three and four indicated that processes such as ABA signalling, leaf senescence, transcription, and translation were all enhanced during chilling stress, but less so after the addition of high light (Fig. 3F, Fig. 5A). The up-regulation of ABA and senescence related processes in response to chilling is expected, as they have been widely reported previously (Capell and Dörffling, 1993, Janowiak, Maas and Dörffling, 2002, Janowiak, Luck and Dörffling, 2003, Masclaux-Daubresse et al., 2007, Wingler and Roitsch, 2008). A reduction in ABA responses upon the addition of high light to the chilling treatment indicates a less stressed state and could result in the re-opening of stomata to allow for continued transpiration and uptake of CO_2_. The reduced induction of senescence related genes upon the addition of high light could also be a significant mechanism behind the mitigating effect of high light during chilling. Delayed leaf senescence and aging allows for enhanced photosynthesis and decreased ROS and has been linked with improved abiotic stress resistance (reviewed by Zhao, Zhao and Wang, 2022). The significant increase in the expression of genes associated with gene expression was somewhat unexpected and counterintuitive. Generally, stress conditions limit energetically consumptive processes such transcription and translation (Muñoz and Castellano, 2012) and translation rates in plants are proportional to temperature (Guillaume-Schöpfer et al., 2020). However, the stress-induced decrease of global translation is often accompanied by a switch to selective transcription and translation of proteins needed for survival during the stress (Holcik and Sonenberg, 2005, Muñoz and Castellano, 2012), in the case of chilling stress, these could be genes involved in processes such as NPQ, chlorophyll metabolism or ABA signalling.

The mechanisms for how high light might mitigate chilling stress suggested here corroborate those reported by Szalai et al. (2018) who found that exposing maize plants to high light (relative to intensity during growth) prior to a severe chilling stress enhanced acclimation to the chilling by modulating genes involved in photosynthesis, chlorophyll biosynthesis, ribosome biogenesis and translation. Thus, it appears that high light has a fundamental impact on plant photosynthesis, metabolism and signalling during chilling stress responses.

### Transcription factors which modulate chilling stress responses and mitigation by high light have been identified

The global transcriptome analyses above identified groups of genes and physiological responses that are significantly impacted during chilling and show a diminished response in the combination with high light. TFs are master regulators of chilling stress responses and can be manipulated to alter tolerance (reviewed: Abdullah, Azzeme and Yousefi, 2022, Sharma et al., 2020, Wang et al., 2016). To identify TFs with the potential to be key drivers of the transcriptional changes observed in modules one, two, three and four, we filtered for TFs which were present in the modules of interest (Table 1 and 2), had enriched binding motifs in the promoters of genes in the same module (Table 3 and 4) and had high kME values for the module in which they belonged (indicating a high level of centrality or connectivity with the other genes in the module). This filtering was based on the premise that TFs and their target genes are co-expressed, such that a change in abundance of a TF alters the interaction of that TF with cis-acting elements in target gene promoters to control stress-responsive gene clusters. Five individual TFs stood out from this analysis: *bHLH99* and *HSF4* from module two and *TCP20, TCP21* and *EREB52* from module three (Fig. 6 and Table 5). All of these are part of TF families often reported in association with stress tolerance and may have promise as candidates for improving chilling tolerance, potentially more so in high light environments. *bHLH99* belongs in the large group of bHLH TFs which have been linked with modulating cold stress responses in several plant species, including maize (reviewed by Guo et al., 2021, Sun, Wang and Sui, 2018). The expression of several bHLH genes was reported as differentially expressed in response to cold in the high-vigour maize variety, Zhongdi175 (Li et al., 2021). Overexpression of the maize bHLH TF, *ZmPTF1*, improved the drought tolerance of transgenic maize, indicating a role for this group of TFs in manipulating abiotic stress responses (Li et al., 2019). As the name indicates, TFs of the heat shock factor (HSF) family often respond strongly to temperature. A study which surveyed several independent maize cold response transcriptomes to detect key genes involved in the response identified *HSF4* as consistently differentially expressed in response to chilling in multiple datasets (Sowiński et al., 2020), illustrating a robust and conserved role for this gene. *TCP20* and *TCP21* form part of the TEOSINTE BRANCHED 1, CYCLOIDEA, PCF1 (TCP) transcription factor family. TCP transcription factors are key modulators of plant growth processes such as branching, floral symmetry and cell cycle and control these processes in response to diverse environmental cues, including abiotic stress (Danisman, 2016). In rice, overexpression of *OsTCP19* improved tolerance to NaCl and mannitol stresses (Mukhopadhyay and Tyagi, 2015), whilst down- regulation of *OsTCP21* increased plant tolerance to cold stress (Wang et al., 2014). Overexpression of maize *TCP42* in Arabidopsis resulted in enhanced drought tolerance (Ding et al., 2019). *EREB52* belongs to the large AP2-EREB transcription factor family which is extensively associated with plant cold stress responses (Ritonga et al., 2021, Zhou et al., 2022b). The most well-known members of this family are the DREBs which form part of the C-repeat binding factor (CBF)-regulon (Liu et al., 2019) which increase the expression of cold-regulated genes and enhance chilling tolerance in maize (Qin et al., 2004, Wang and Dong, 2009, Wang, Yang and Yang, 2011). Thus, the five TFs that arose from analyses presented here are clearly implicated to impact maize chilling responses and tolerance and warrant further exploration.

### F7 is more chilling tolerant than B73

Whilst significant differences were observed between F7 and B73 for some of the physiological parameters measured here (Fig. 7), there were no significant interactions between accession and treatment, indicating a lack of accession-specific differential response to the stress that would indicate variation in chilling tolerance. However, this may be due to the relatively short (15 hour) chilling treatment period imposed here rather than a lack of difference in chilling tolerance between the accessions. Enders et al. (2018) reported that maize grown under a 24 hour period of chilling had few changes in growth inhibition (plant height, area, and width) compared with plants under control conditions. Thus, observing physiological differences in the stress response between accessions may require a longer treatment period. There were, however, transcriptomic indicators which suggested greater chilling tolerance in F7. Firstly, F7 had substantially more DEGs than B73 in response to chilling and chilling plus high light (Fig. 3D) which may suggest a stronger, more effective response with more genes responding to the stress. The same type of result was reported by Zhou et al. (2022a) when they compared B73 with the chilling-tolerant Mo17 accession and Li et al. (2019) when comparing chilling-tolerant M54 and chilling-sensitive 753F accessions. Secondly, individual genes which have previously been reported to confer chilling tolerance showed different patterns of expression in F7 compared with B73 (Fig. 2). The expression of *ZEP2* was significantly down-regulated in B73 in response to chilling and chilling plus high light whereas the expression is up-regulated in F7 in these treatments (Fig. 2A). Conversely, *VDE3b* was up-regulated in B73 but down-regulated in F7 in response to chilling and chilling plus high light (Fig. 2B). These results suggest a greater accumulation of zeaxanthin in B73, which is negatively associated with chilling tolerance (Fracheboud et al., 2002). Another such example of chilling tolerance in F7 is the differences in expression of *CRR1* and *MPK8*. *CRR1*, a positive regulator of chilling stress (Zeng et al., 2021), was up-regulated in F7 and down regulated in B73 whereas *MPK8*, a negative regulator of chilling stress in the same signalling pathway (Zeng et al., 2021), showed the opposite expression (Fig. 2C and D). These are some examples of possible candidates for conferring chilling tolerance, but it would be beneficial to perform the same type of transcriptome analysis on plants exposed to longer periods of chilling stress when the physiological differences between accessions may become more apparent. Interestingly, comparison of the F7-specific chilling-responsive transcriptome (i.e. DEGs in F7, but not in B73) with that of another relatively chilling-tolerant accession, Mo17 (published by Zhou et al., 2022a) suggested common mechanisms such as brassinosteroid modulated signalling might help to enhance tolerance. Brassinosteroid application to chilling-sensitive species such as rice, tomato, cucumber, Arabidopsis and maize improves plant performance by promoting the expression of *COR* genes, master regulators of the cold response as well as mechanisms including antioxidant mobilization, membrane fluidity and Ca^2+^ influx (reviewed by Ramirez and Poppenberger, 2020).

To conclude, the work here provides novel insights into the role of high light during plant exposure to chilling conditions. The addition of high light was able to alleviate the negative impact of chilling on photosynthesis, photoinhibition and membrane integrity induced by chilling. Additionally, of the ±2500 annotated transcription factor genes in maize (Lin et al., 2014), five were identified as strong candidates in driving the mitigating changes in transcription brought about by high light in combination with chilling. Importantly the transcriptional responses associated with improved chilling tolerance of the F7 accession are very similar to those reported for other accessions and so we propose these reflect a common transcriptional response in maize accessions with enhanced chilling tolerance.

## Materials and methods

### Plant material and growth conditions

Seeds of the maize (*Zea mays*) inbred lines B73 and F7 were obtained from the International Rice Research Institute (IRRI) and the U.S. National Plant Germplasm System (NPGS) respectively. Seeds were germinated on wet filter paper in the dark at 28°C for three days after which each germinated seed was transferred into a two litre pot containing a mixture of 2 parts nutrient-rich compost (Levington Advance M3, ICL, Ipswich, UK) to 1 part topsoil (Westland, Dungannon, Northern Ireland), 10 ml Miracle-Gro all-purpose fertiliser beads and 15 ml Miracle-Gro magnesium salt (Scotts Miracle-Gro, Marysville, OH, USA) and placed in a growth room (Conviron Ltd, Winnipeg, CA) set at 28/20°C day/night, 65% relative humidity, 400 μmol photons m^-2^ s^-1^ and a 12 hour photoperiod. 15 days post germination, the plants were transferred to a growth cabinet (Percival E-41HO, CLF Plant Climatics GmbH, Wertingen, Germany) set to the same conditions as the growth room. Plants were grown in three batches to which different experimental treatments were applied (see below). Each batch contained eight plants per accession, of which four plants per accession were randomly assigned to the experimental control and four plants per accession were exposed to the experimental treatment.

### Experimental treatments and sampling

Three discrete experiments were performed within a four-week period, each with a different treatment applied to a separate batch of plants. For each of the three experiments four control plants per accession were harvested three hours after the start of the photoperiod on day 20 (post germination). At the end of day 20, the remaining four plants in the cabinet were treated with one of three treatments (i.e. treatments occurred on night 20 and three hours into the photoperiod of day 21 post germination). A chilling treatment was applied to the first batch of plants, with a temperature of 8/5°C (D/N). Plants were harvested following 12 hours chilling night (5°C) and three hours chilling day (8°C). A high light treatment of 1000 µmol photons m^-2^ s^-1^ during the three hours day was applied to the second batch of plants. This treatment was applied at the beginning of the photoperiod on day 21 (post germination) and plants were harvested following three hours high light. Finally a combined chilling and high light treatment (8/5°C; 1000 µmol photons m^-2^ s^-1^) was applied to the third batch of plants and plants were harvested following 12 hours chilling night and three hours combined chilling and high light day. All three independent experiments were conducted in the same growth cabinet.

Tissue harvesting was performed three hours into the photoperiod on day 20 (control plants) or day 21 (treated plants). All samples were taken from expanded leaf three (the third leaf to emerge), before expansion of leaf four. For all plants, two rectangular leaf cuttings (avoiding the midrib) were obtained and flash-frozen in liquid nitrogen for subsequent RNA extraction. Additionally, discs of 6 mm diameter were punched from each leaf for subsequent physiological measurements:

For the electrolyte leakage assay (relative injury), three discs per leaf were 200μL MilliQ water for a 30 minute wash period to remove electrolytes resulting from the punching of the disc. Each disc was then placed into 1 mL MilliQ water and left to soak for 20 hours.

For the chlorophyll fluorescence assay, three discs per leaf were each placed adaxial side facing down into the well of a 96-well plate and held in place with a small piece of sponge dampened in MilliQ water to prevent dehydration.

The MDA and chlorophyll assays were determined on the same sample. Three discs per leaf were pooled and flash-frozen in liquid nitrogen and stored at - 80°C for subsequent analysis.

### Relative injury measurement

Following the 30 minute wash period and 20 hours soaking, samples were shaken at 300 rpm for 15 minutes (Thermomixer-Mixer HC, Starlab (UK) Ltd, Milton Keynes, UK). The electrical conductivity of a 100 µl aliquot of the resultant solution was measured (LAQUAtwin EC-11 conductivity meter, Horiba Instruments, Kyoto, Japan). Following this measurement (Time 1), samples were heated at 99°C for three hours and cooled to room temperature. Each sample was vortexed and electrical conductivity was re-measured (Time 2). For each measurement of conductivity, samples were measured at least twice (a new aliquot was taken each time) until the reading was stable between aliquots. A blank (MilliQ water) was also measured at each time point.

Relative injury of the leaf material was calculated according to the following formula (Thalhammer et al., 2020):

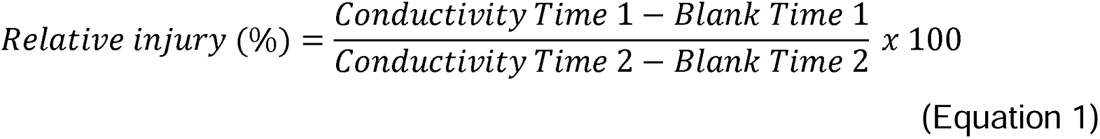

### Chlorophyll fluorescence measurement

Chlorophyll fluorescence was used to measure F_v_/F_m_ and NPQ (Fracheboud et al., . 1999; Baker et al., 2008). The leaf discs harvested into 96-well plates were immediately dark adapted for one hour. For chilling treatments (both chilling and the combined chilling and high light treatment) this dark adaptation was carried out in chilling conditions. For the high light and control treatments this dark adaptation was carried out at room temperature. Following dark adaptation, chlorophyll fluorescence was measured using a FluorCam (Photon Systems Instruments, Drásov, Czech Republic). Leaf discs were exposed to a 10 minute light period followed by a 12 minute dark relaxation period. F_v_/F_m_ was measured on dark-adapted leaves at the start of the measurement period. NPQ was obtained from saturating pulses in the light and dark. In the light, saturating pulses were given every 20 seconds for the first minute, then once per minute. In the dark, saturating pulses were given every 20 seconds for the first minute, then once per minute for two minutes, then once every three minutes for the remaining nine minutes. For samples resulting from the chilling treatments (the chilling treatment and the combined chilling and high light treatment) the FluorCam measurement was carried out at a leaf temperature of 8°C, achieved by placing a cold plate inside the FluorCam (Solid State Cold/Hot Plate AHP- 301CPV, ThermoElectric Cooling America Corporation, Chicago, IL, USA). For samples resulting from the high light treatment the FluorCam measurement was carried out at a leaf temperature of 20°C.

### Biochemical assays

Samples harvested for MDA and chlorophyll measurements were cryo-ground (TissueLyser II, Qiagen, Hilden, Germany). Glass beads were inserted into sampling tubes prior to harvest to expedite the grinding process.

### MDA and chlorophyll quantification

Each set of three discs harvested for MDA and chlorophyll assays was extracted in 1ml 80% ethanol (v/v) at 4°C for 72 hours. Samples were then vortexed (15 s) and centrifuged (5 mins, 4°C, 13,000rpm).

For the chlorophyll measurement, a 150 μl aliquot of the ethanol extract was added to a 96-well plate in duplicate, along with an ethanol blank, and absorbance was measured at 632 nm, 649 nm, 665 nm and 696 nm (Synergy HT Plate Reader, BioTek, Winooski, VT, USA). Chlorophyll was determined according to Ritchie et al. (2008).

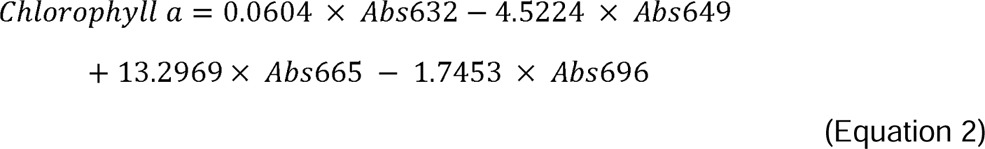

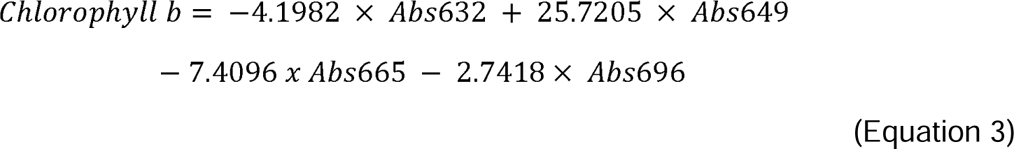

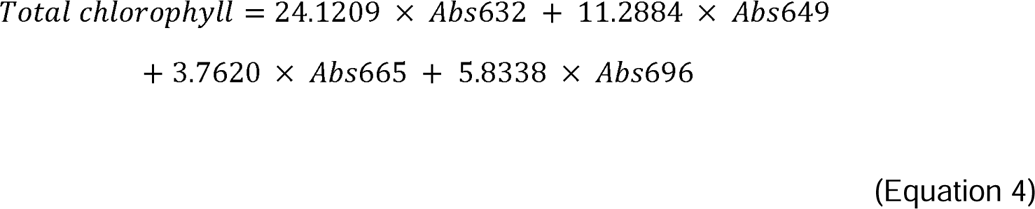

For the MDA measurement, two 250 μl aliquots were taken from each ethanol extract. One aliquot was added to 250 μL “Minus TBA” solution and one aliquot was added to 250 μL “Plus TBA” solution. The “Minus TBA” solution contained 200 g/L trichloroacetic acid and 0.1 g/L butylated hydroxytoluene. The “Plus TBA solution” consisted of “Minus TBA” with the addition of 6.5 g/L thiobarbituric acid. Samples with plus/minus TBA solution were vortexed (15 s), heated (25 mins, 95°C), cooled to room temperature then centrifuged (5 mins, 4°C, 13,000rpm). A 200 μl aliquot was then added to a 96-well plate in duplicate, along with a blank (“Plus TBA” in 80% ethanol). Absorbance was measured at 440 nm, 532 nm and 600 nm (Synergy HT Plate Reader, BioTek, Winooski, VT, USA). MDA was determined according to the method of Hodges et al. (1999), using the corrected equation by Landi (2017).

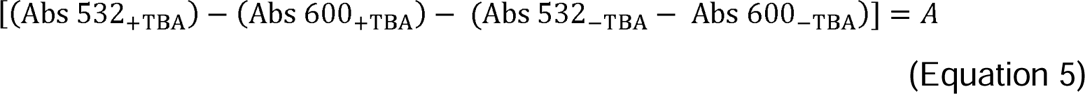

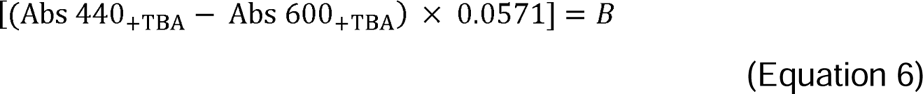

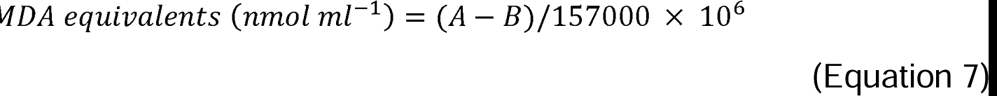

### Total RNA extraction, cDNA library preparation and Transcriptome Sequencing

Total RNA was extracted from leaf samples of the control and treated (chilling, high light and chilling plus high light) plants using the RNeasy Plant Mini Kit (Qiagen, Germany) according to the manufacturer’s instructions. Genomic DNA contamination was removed from each RNA sample using the RNase-Free DNase Set (Qiagen, Germany). RNA degradation and contamination were checked on 1% agarose gels, then RNA quality and concentration were determined with the RNA600 Pico Assay using the Agilent 2100 Bioanalyzer (Agilent Technologies, USA) and ND-200 nanodrop (NanoDrop Technologies Inc., USA). For each sample, a cDNA library was prepared from 200 ng RNA starting material using the QuantSeq 3’ mRNA-Seq Library Prep kit FWD for Illumina (Lexogen, Austria) according to the manufacturer’s instructions. Library quality and concentration was determined with the High Sensitivity DNA assay using the Agilent 2100 Bioanalyzer (Agilent Technologies, USA) and Qubit® dsDNA HS Assay Kit (Invitrogen, USA). Finally, the libraries were sent to the Novogene Cambridge Genomic Sequencing Centre for sequencing using the Illumina NovaSeq 600 PE150 sequencing platform and strategy. At least 6 Gb of raw data per sample was generated for subsequent transcriptome analysis (refer to Supp. dataset 9 for the number of raw reads generated for each sample).

### Transcriptome data processing

Due to the nature of the library preparation kit used only the forward sequencing reads were used for data analysis. The quality of raw sequencing data (.fastq format) was assessed and controlled using the FastQC platform (version 0.11.4, Andrews, S. 2010, http://www.bioinformatics.babraham.ac.uk/projects/fastqc/). Adapter trimming and filtering of all low quality reads was performed using BBDuk (https://www.geneious.com/plugins/bbduk/) with the following parameters: k=13, ktrim=r, useshortkmers=t, mink=5, qtrim=r, trimq=20, minlength=50. The Zm-B73- REFERENCE-NAM-5.0 transcriptome was downloaded from MaizeGDB (https://www.maizegdb.org/genome/assembly/Zm-B73-REFERENCE-NAM-5.0) and used to build a Salmon reference index which was used to quantify the cleaned reads (Salmon version 1.5.2). For Salmon quantification (Patro et al., 2017), all parameters were left as default, except the --noLengthCorrection parameter was used. The Salmon alignment and quantification results were checked using MultiQC (https://multiqc.info/). Refer to Supp. dataset 9 for the number of reads remaining after BBDuk trimming and the % alignment for reads to the B73 reference genome. To check that biological replicates clustered together and to visualise how the treatments differed from controls, a principal component analysis (PCA) was performed using the ggfortify package in R (https://cran.r-project.org/web/packages/ggfortify/index.html). For this, the transcripts per million (TPM) outputs from the Salmon quantification were used. To filter out very lowly expressed genes, only genes with a total of 36 TPMs across all 48 samples were included. The filtered data were then normalised to account for library size and transformed to a log scale using variance stabilising transformation (vst) to allow for easier visualisation.

### Identification of differentially expressed genes (DEGs), weighted Gene Correlation Network Analysis (WGCNA) and functional annotation of clusters and

To determine the changes in expression of individual genes, the DESeq2 package in R was used (version 4.2, Love, Huber and Anders, 2014, https://bioconductor.org/packages/release/bioc/html/DESeq2.html). The quant.sf file generated for each sample from the Salmon quantification was used as the input for the DESeq2 analysis. Independent DESeq2 analyses were performed for the three independent experiments (i.e. samples from each treatment were compared with the corresponding control for that experiment).

The WGCNA package in R (version 1.71, Langfelder and Horvath, 2008, https://cran.r-project.org/web/packages/WGCNA/index.html) was used to cluster genes which had similar patterns of expression. For the analysis the TPM outputs from the Salmon quantification were used. To filter out genes with very low expression levels, only genes with a total of 36 TPMs across all 48 samples were included. The WGCNA one-step network construction and module detection function were used to construct the gene network and identify modules using a “signed” network type and minimum module size of 30 genes. All other parameters were left as default. The lists of genes within each module and module eigengene values were then extracted for downstream analysis.

To gain further biological insight into modules with interesting expression patterns, Gene Ontology (GO) enrichment analyses were performed using AgriGO v2 (Tian et al., 2017, http://systemsbiology.cau.edu.cn/agriGOv2/) with the Maize AGPv4 (Maize-GAMER) genome set as the background reference.

### Determination of transcription factors in gene lists and enriched transcription factor binding sites to identify candidate transcription factors

The transcription factors present in the combined module gene lists were identified using the Plant Transcription Factor and Protein Kinase Identifier and Classifier (iTAK) tool (Zheng et al., 2016, http://itak.feilab.net/cgi-bin/itak/index.cgi). For this, the unspliced gene sequences for each gene were downloaded from Biomart (https://plants.ensembl.org/biomart/martview/) using the Ensembl Plants Gene 54 database and Zm-B73-REFERENCE-NAM-5.0 dataset. To identify the enriched transcription factor binding sites in the promoters of genes in the combined module gene lists, a Simple Enrichment Analysis (SEA) using the MEME Suite of Motif-based sequence analysis tools (Bailey and Grant, 2021, https://meme-suite.org/meme/index.html) was performed. The promoter region 1000 bp upstream of the transcription start site of each gene was downloaded from Biomart. This was compared to a manually curated list of control promoter sequences which were from a list of genes which showed no significant differential expression in any of the treatments (i.e. *p*-adj > 0.05 in chilling, chilling plus high light and high light treatments). The CIS-BP 2.00 Single Species DNA motif database for *Zea mays* was used to test for enrichment. To identify transcription factor candidates, the top 18-20 enriched TF binding motifs from the SEA were scanned to determine whether they were present in the list of TFs in the iTAK output. Transcription factors identified were then filtered for a module membership (kME) value greater than 0.9. The kME values indicate the centrality of a gene within a WGCNA module.

## Statistics

The adjusted *p*-values from the DESeq2 output were used to identify statistically significantly differentially expressed genes. DESeq2 uses a Wald test for significance testing, and Benjamini and Hochberg to adjust *p*-values for multiple comparisons. Two-way ANOVA was performed to identify differences in relative injury and chlorophyll fluorescence parameters, testing for effects of accession, treatment, and an interaction between accession and treatment. All statistics were performed using R (R Core Team, 2022).

## Supplemental data

- Supplemental dataset 1: DESeq2 data for genes in figure 2, 4, 5 and 6 (excel doc – sheet 1).
- Supplemental dataset 2: TPM data for genes in figure 2, 4, 5 and 6 (excel doc – sheet 2).
- Supplemental dataset 3: List of DEGs common in F7 and Mo17, but not B73, in response to chilling (excel doc – sheet 3).
- Supplemental dataset 4: Lists of genes present in each WGCNA module (excel doc – sheet 4).
- Supplemental dataset 5: Results from GO enrichment analyses of WGCNA modules of interest (excel doc – sheet 5).
- Supplemental dataset 6: Comparison of enriched GO terms present in modules one versus two and modules three versus four (excel doc – sheet 6).
- Supplemental dataset 7: List of all TFs present in modules one, two, three and four (excel doc – sheet 7).
- Supplemental dataset 8: Transcription factor binding motifs enriched in the promoters of genes in modules one, two, three and four identified using the MEME Suite of Motif-based sequence SEA tool (excel doc – sheet 8).
- Supplemental dataset 9: Supp. dataset 9: Illumina sequencing output (raw reads) for each sample, reads remaining after BBDuk trimming and the % of these reads which aligned to the B73 reference genome using Salmon alignment (excel doc – sheet 9).

## Author contributions

LC, ACB and JK conceived and designed the experiment with input from JR and JMH, LC, ACB and JR performed the measurements. LC and ACB analyzed the data. LC and ACB wrote the manuscript with contributions from all authors.

