## Supplementary material for "High light can alleviate chilling stress in maize": Supp. figure

**Supplemental figures**

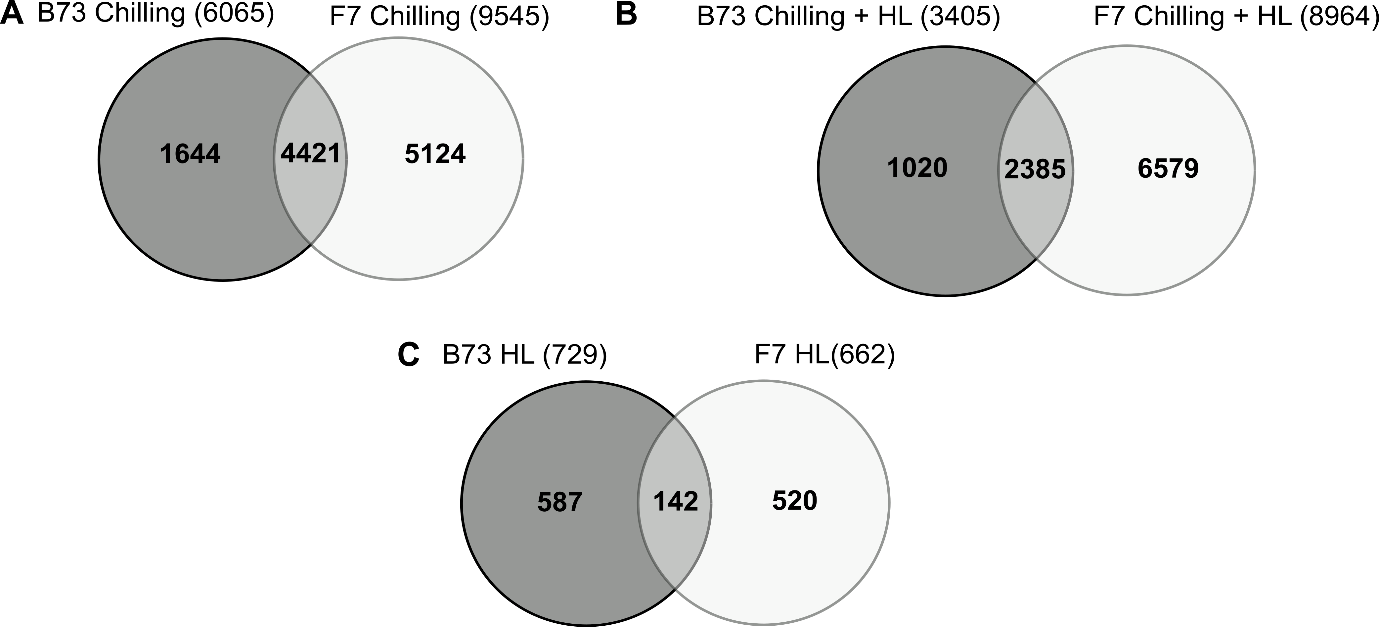

**Supplemental Figure 1: Venn diagram showing the extent of overlap between significantly differentially expressed genes (DEGs) in each accessions in each treatment.** **A**, overlap of DEGs between B73 and F7 in response to chilling treatment. **B**, overlap of DEGs between B73 and F7 in response to chilling and high light (HL) treatment. **C**, overlap of DEGs between B73 and F7 in response to high light (HL) treatment.

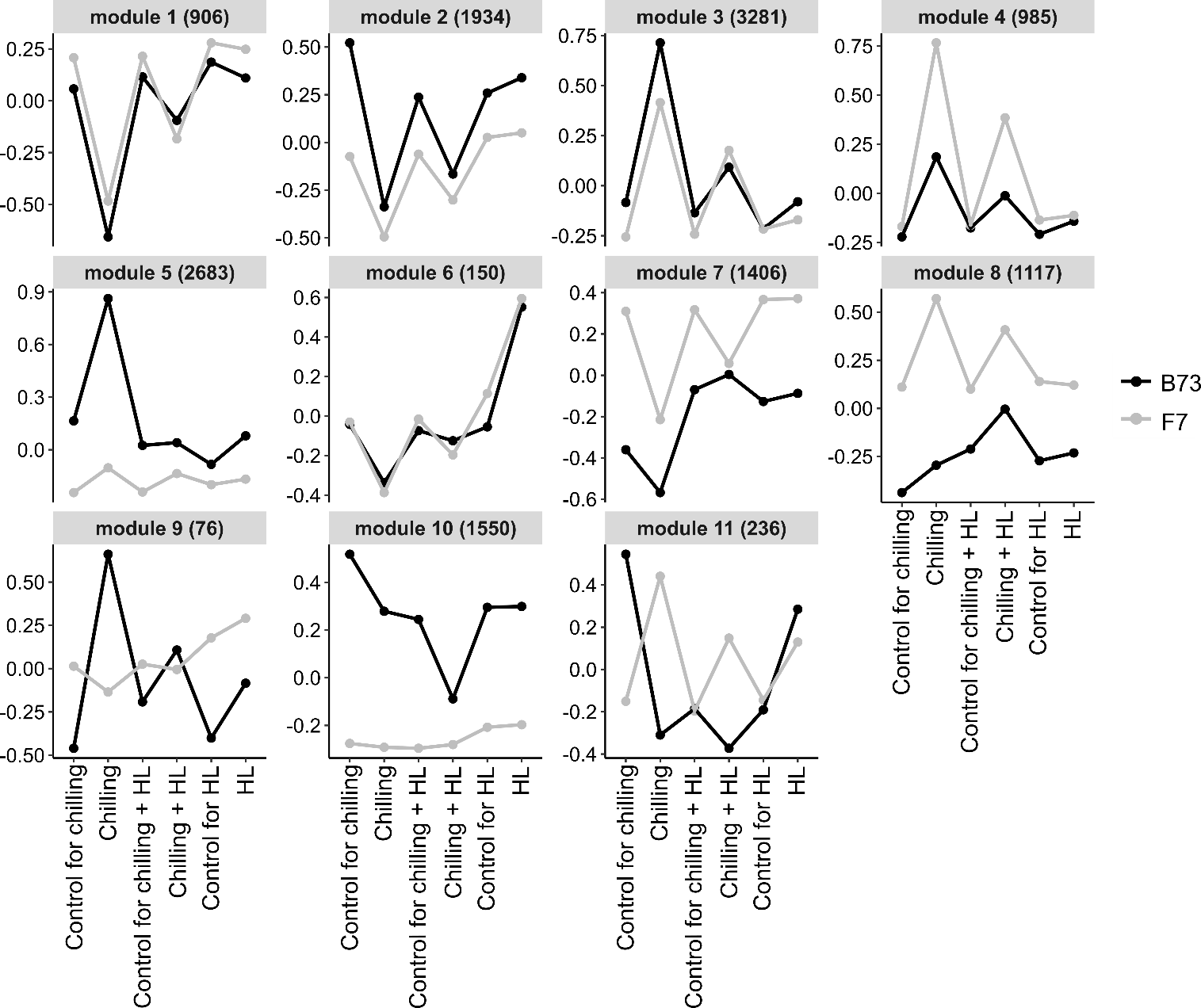

**Supplemental Figure 2: Eigengene expression of the eleven modules identified from a WGCNA performed on the transcriptome data.** Filtered TPM data was used for a WGCNA to identify modules of genes with similar patterns of expression. All plots show eigengene values representing the gene expression profile in each module. Numbers in brackets represent the number of genes in each of the modules. Lists of genes in each module can be found in Supp. dataset 4.

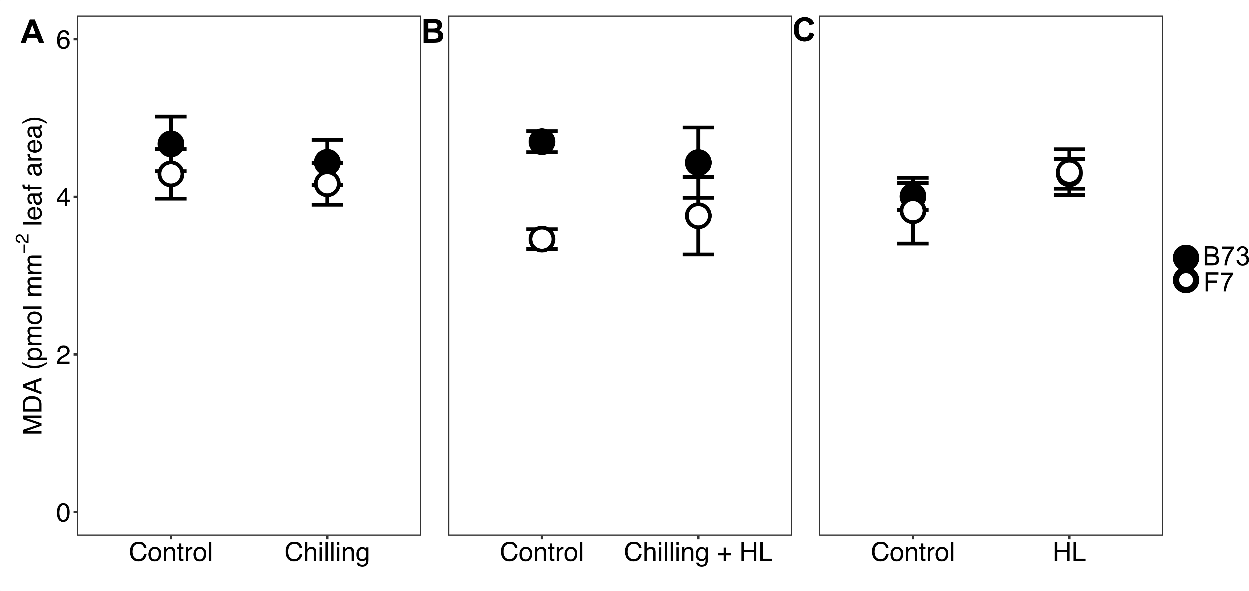

**Supplemental Figure 3:** Malondialdehyde (MDA) content in leaves of maize plants exposed to chilling (**A**); chilling plus high light (HL, **B**); or **C**, high light (**C**).

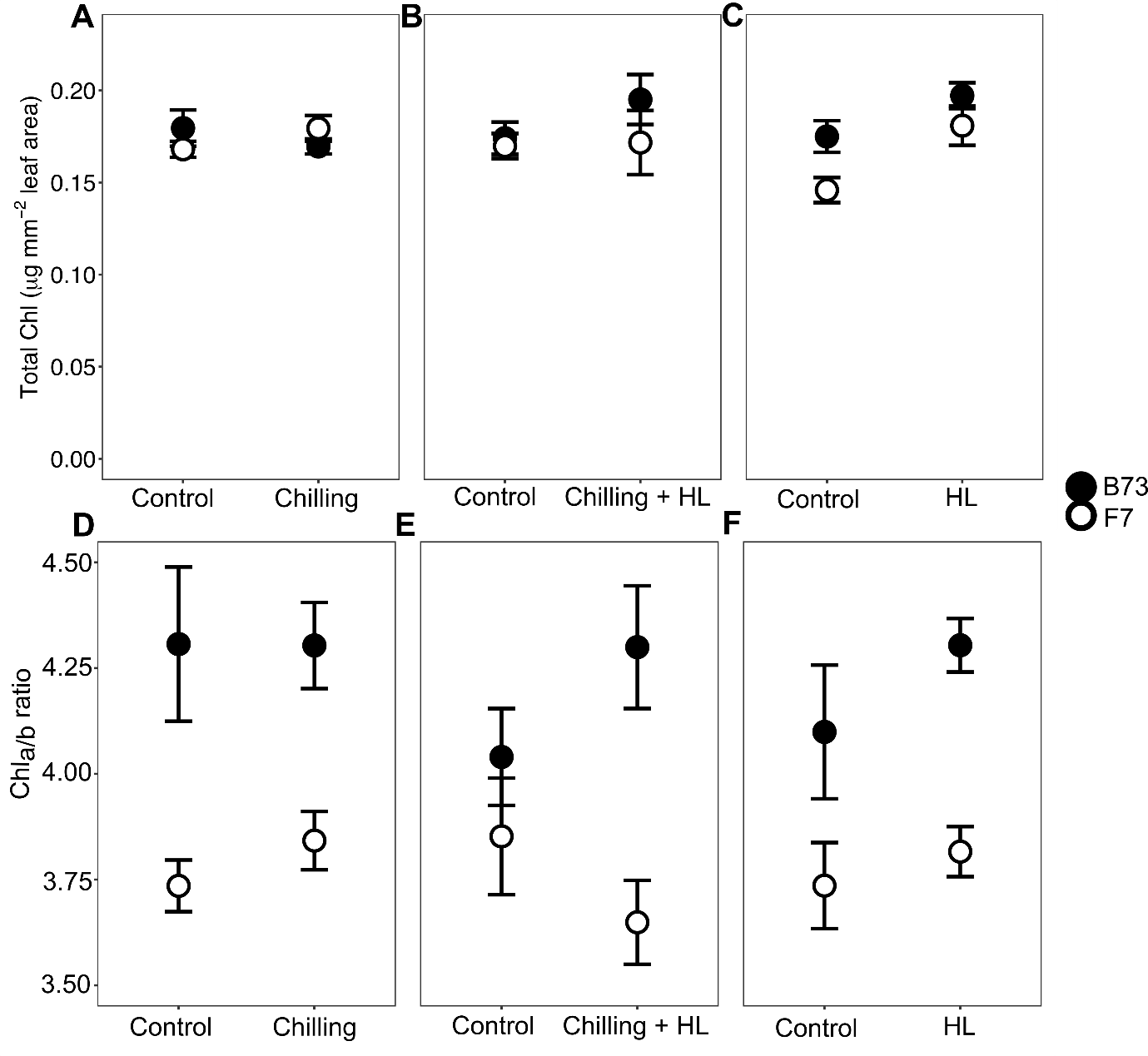

**Supplemental Figure 4:**  Total chlorophyll content and chlorophyll a/b ratio in leaves of maize plants exposed to chilling (**A/D**); chilling plus high light (HL, **B/E**); or **C**, high light (**C/F**).
